# Integration of chromatin accessibility and gene expression data with cisREAD reveals a switch from PU.1/SPIB-driven to AP-1-driven gene regulation during B cell activation

**DOI:** 10.1101/2023.01.09.522862

**Authors:** Amber M.L. Emmett, Amel Saadi, Matthew A. Care, Gina M. Doody, Reuben M. Tooze, David R. Westhead

## Abstract

Human B cell differentiation into antibody secreting plasma cells is a critical process in the adaptive immune response, whose regulation at the genetic level remains incompletely understood. To reveal the temporal sequence of transcription factor driven cellular changes we generated chromatin accessibility (ATAC-seq) and gene expression (RNA-seq) data from *in vitro* differentiation of human B cells into plasma cells using a published protocol for differentiation up to the plasma cell stage. Using a new computational method, cisREAD (cis-Regulatory Elements Across Differentiation), we defined a core set of *cis*-regulatory elements that are confidently linked to dynamic transcription factor binding and changes in gene expression across the mature B lineage. Here we describe how cisREAD identifies regulatory element ‘communities’, based on chromatin accessibility and transcription factor co-occupancy, and prioritizes those whose accessibility predicts differential gene expression through regularized regression models. Through downstream analyses of cisREAD-predicted regulation, we show how transcription factors reshape B cell epigenomes and transcriptomes in response to differentiation stimuli. Our results confirm roles for OCT2, IRF4 and PRDM1 in plasma cell differentiation, and reveal that a shift from PU.1/SPIB-driven to AP-1-driven gene regulation is a key determinant of B cell activation.

**GRAPHICAL ABSTRACT:** Integration of epigenomic and transcriptomic datasets with the cisREAD method, followed by clustering and network analysis, reveals that gene regulation shifts from PU.1/SPIB to AP-1 upon B cell activation.

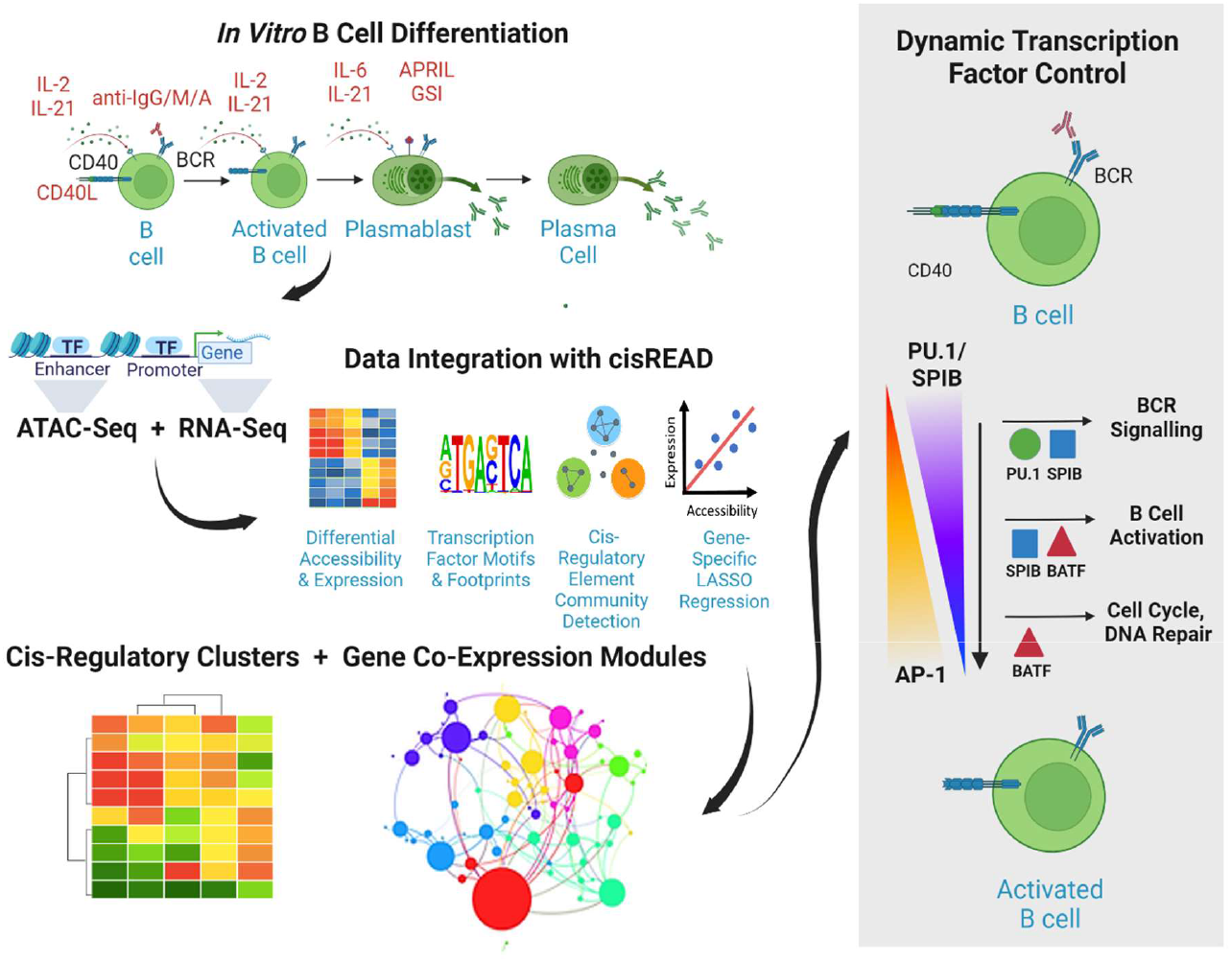

## INTRODUCTION

The differentiation of B cells (BCs) to plasma cells (PCs) is essential to humoral immunity, and is controlled through a dynamic gene regulatory network, shaped by epigenetic reprogramming through transcription factors (TFs). This process is initiated by activation signalling, upon which B cells are stimulated to divide and a subset differentiate into quiescent plasma cells, capable of secreting large quantities of antigen-specific antibodies. As B cells transition through proliferative activated B cell (ABC) and plasmablast (PB) states, regulatory control shifts from a transcriptional circuit upholding B cell identity (including PAX5 and BACH2) to a mutually antagonistic network promoting plasma cell fate (including IRF4, PRDM1 and XBP1) (1, 2). Whilst this collective of transcriptional regulators is well characterised, there is an incomplete understanding of how transcription factors reprogram gene expression in response to activation and differentiation stimuli.

Chromatin accessibility, measurable by the Assay for Transpose Accessible Chromatin (ATAC-seq), acts as a readout for the epigenetic mechanisms, such as histone modifications and DNA methylation, which determine chromatin permissiveness (3). Integrating ATAC-seq data with transcriptomic profiling through RNA-seq, has allowed researchers to map healthy and malignant B cell regulomes in a number of organisms, primary cells and *in vivo* and *in vitro* differentiation systems, leading to new insight into regulation of the B lineage (4–7).

Linking accessible regulatory elements to gene expression remains a challenge. Three dimensional chromatin conformation technologies like Hi-C have been used to determine regulatory chromatin domains in B cells (8–10). Yet tens of millions of cells are required for sufficient resolution to detect enhancer-promoter interactions, as opposed to tens of thousands for ATAC-seq, limiting their use (11, 12). Instead researchers often rely on assigning cis-regulatory elements to their nearest gene, which is not always the case for distal interactions (13). Whilst sophisticated methods exist to assign regulatory elements to genes, they operate in a cell-specific manner and often require many input epigenetic datatypes (14–17). Here we introduce a new method, based on correlating differential cis-regulatory element (CRE) accessibility with differential gene expression, designed to identify and prioritize transcription-factor bound cis-regulatory elements across differentiation (cisREAD). We apply this method to 19 sample-matched ATAC-seq and RNA-seq samples taken from up to three donors at fine-grained intervals across human *in vitro* B cell activation stimulated by CD40L and plasma cell differentiation induced by APRIL (18, 19).

Through our data-driven global analysis of transcriptional regulation we reveal distinct regulatory determinants for a core set of lineage-specifying and signal-inducible transcription factors. Notably, we highlight how a shift from PU.1/SPIB-driven regulation to AP-1 mediated control is central feature of the activated B cell state that precedes commitment to plasma cell fate.

## METHODS

### B cell differentiation

B cell populations were isolated from the peripheral blood of 3 donors using MACS separation (total B-cells for time-points to day 3, and memory B-cell-enriched for time points after day 6 and 13) and long-lived plasma cells were generated *in vitro* following published protocols (Figure 1A) as previously described (18, 20). Briefly, B cells were exposed to activating conditions including F(ab’)2 anti-IgG/A/M, IL-2, IL-21 and irradiated CD40L L-cells at day 0 to stimulate B cell activation. Cells were sampled by careful removal from the stromal cell layer at indicated time points. Cells were transferred at day 3 to conditions with cytokines IL-2 and IL-21 alone, and plasmablasts at day 6 were driven towards a long-lived plasma cell phenotype through further cytokine signalling (IL-21, IL-6 and APRIL) and the addition of γ-Secretase Inhibitor (GSI) (19). ATAC-seq and RNA-seq experiments, measuring chromatin accessibility and gene expression, were performed at 9 time-points across the *in vitro* differentiation process, yielding a dataset of 19 samples (Figure 1B).

**Figure 1.**
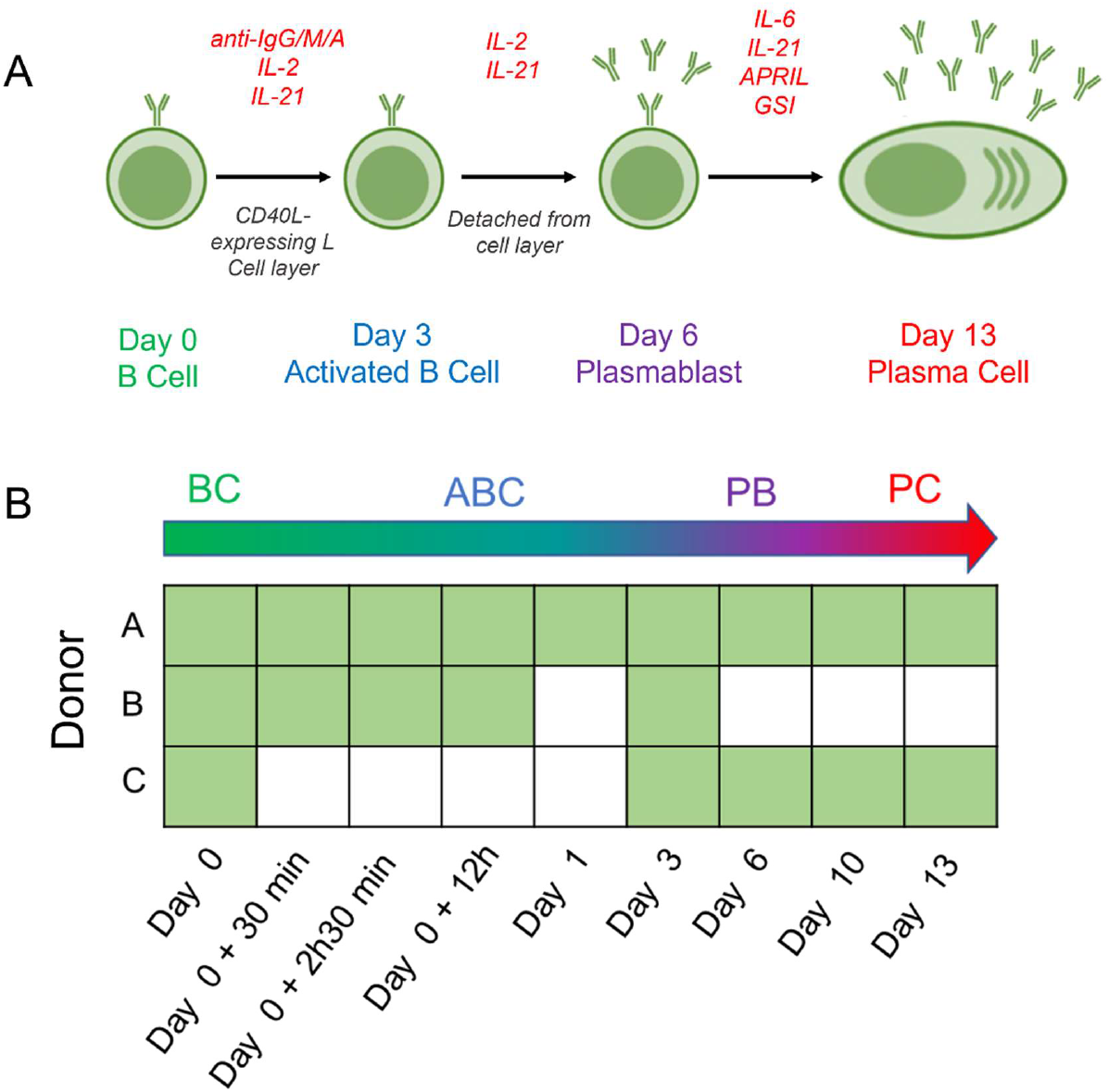
*In vitro* system of human B cell activation plasma cell differentiation. A) B cells are first extracted from the peripheral blood of donors and are cultured and activated in the presence of the CD40 ligand, anti-Ig and IL-2 and IL-21 cytokines. Activated B cells then undergo plasma cell differentiation, driven through removal of CD40L and anti-IgG/M/A and are pushed towards a mature plasma cell phenotype through removal of IL-2 and provision of APRIL, γ-Secretase inhibitor (GSI) and IL-6. B) Donor (A, B, C) matched ATAC-seq and RNA-seq datasets taken at nine time-points across in vitro plasma cell differentiation, from 1-3 biological replicates, yielding a total of 19 samples. BC, B cell; ABC, activated B cell; PB, plasmablast, PC; plasma cell.

### ATAC-seq & RNA-seq

ATAC-seq was performed according to the protocol described by Buenrostro et *al.* (21). Briefly, cells were harvested at indicated time points with 50,000 cells used per time point. Cells washed once with 50 μl of cold 1x PBS buffer were re-suspended in 50 μl of cold lysis buffer (10 mM Tris-HCl, pH 7.4, 10 mM NaCl, 3 mM MgCl2, 0.1% IGEPAL CA-630). Transposition reactions were performed at 37°C for 90 min using Tn5 Transposase (Illumina Cat #FC-121-1030) in 2x TD Buffer (Illumina Cat #FC-121-1030) and nuclease free H2O. Immediately following transposition, DNA was purified using a Qiagen MinElute Kit with elution into 10mM Tris buffer, pH 8. Initial amplification of transposed fragments was performed using customized Nextera PCR Primer 1* and customised Nextera PCR Primer 2* with NEBNext High-Fidelity PCR mix (New England Labs Cat #M0541) with 8-14 (median 12) cycles of amplification. Amplification was monitored using qPCR and Bioanalyser (Bioanalyzer High Sensitivity DNA kit). Amplified libraries were cleaned up using Qiagen PCR Cleanup Kit and eluted in 10mM Tris Buffer, pH 8 and quantified using KAPA Library Quant Kit for Illumina Sequencing Platforms (KAPABiosystems). Samples were sequenced on a NextSeq500 with 2×75-bp paired-end reads.

At matched time points shown in Figure 1, samples were removed and total RNA was extracted using RNeasy Plus Micro Kit (QIAGEN) subjected to DNAseI treatment (DNA Free; Ambion), and subsequently amplified using the Illumina TruSeq Stranded Total RNA Human/Mouse/Rat kit (Illumina). Libraries were sequenced on the NextSeq500 (Illumina), using 76-bp single-end sequencing (NextSeq500).

### Data Processing

ATAC-seq and RNA-seq FASTQ files were first quality checked with FastQC. Trim Galore was then used with default settings to remove Illumina sequencing adaptors and low-quality reads (Q < 20).

Trimmed, paired ATAC-seq reads were aligned to the human genome (NCBI GRCh38 decoy version) using bowtie2 (--very-sensitive). Post-alignment, low-quality mappings (MAPQ < 20) were filtered out with Samtools and duplicates were removed with Picard tools. Bedtools was also used to filter out ENCODE and mitochondrial blacklists (22), and select fragments <100bp to isolate nucleosome-free regions. Coordinates were shifted with deepTools +4 bp on the positive strand and −5 bp on the negative to centre on the Tn5 cutting site. Narrow peaks were called with Macs2 in paired-end mode with q < 0.05. Consensus peak sets were constructed using the DiffBind R package to retain peaks present in at least 50% of donors for each time-point. A count matrix was then produced from the union of all consensus peaks, across all samples, and accessibility signal was normalised using DESeq2s median of ratio’s method.

Trimmed RNA-seq reads were mapped against the GRCh38 decoy-aware human transcriptome and quantified with Salmon, correcting for GC bias and sequence bias. Transcript counts were aggregated to gene level with the tximport R package and normalised using DESeq2’s median of ratios method.

ATAC-seq and RNA-seq count matrices were *vst* transformed within DESeq2 and subject to preliminary analysis in R. Principal components were computed and a PCA biplot was produced with the ggplot2 package. Euclidean sample distances were hierarchically clustered and visualised with the pheatmap package. These unsupervised analyses revealed that samples cluster tightly by differentiation stage, with no evidence of batch-effects or outliers (Figure S1).

### cisREAD Methodology

**W**e introduce a computational method designed to identify and prioritise cis-Regulatory Elements Across Differentiation (cisREAD) from chromatin accessibility (ATAC-seq) and gene expression (RNA-seq) datasets spanning a cellular lineage. In this section we outline the cisREAD approach, which uses community detection and LASSO regression to identify gene-specific cis-regulatory elements, targeted by core transcription factors. Our approach is represented graphically in Figure 2, and the software is publicly available as an R package at https://github.com/AmberEmmett/cisREAD.

**Figure 2.**
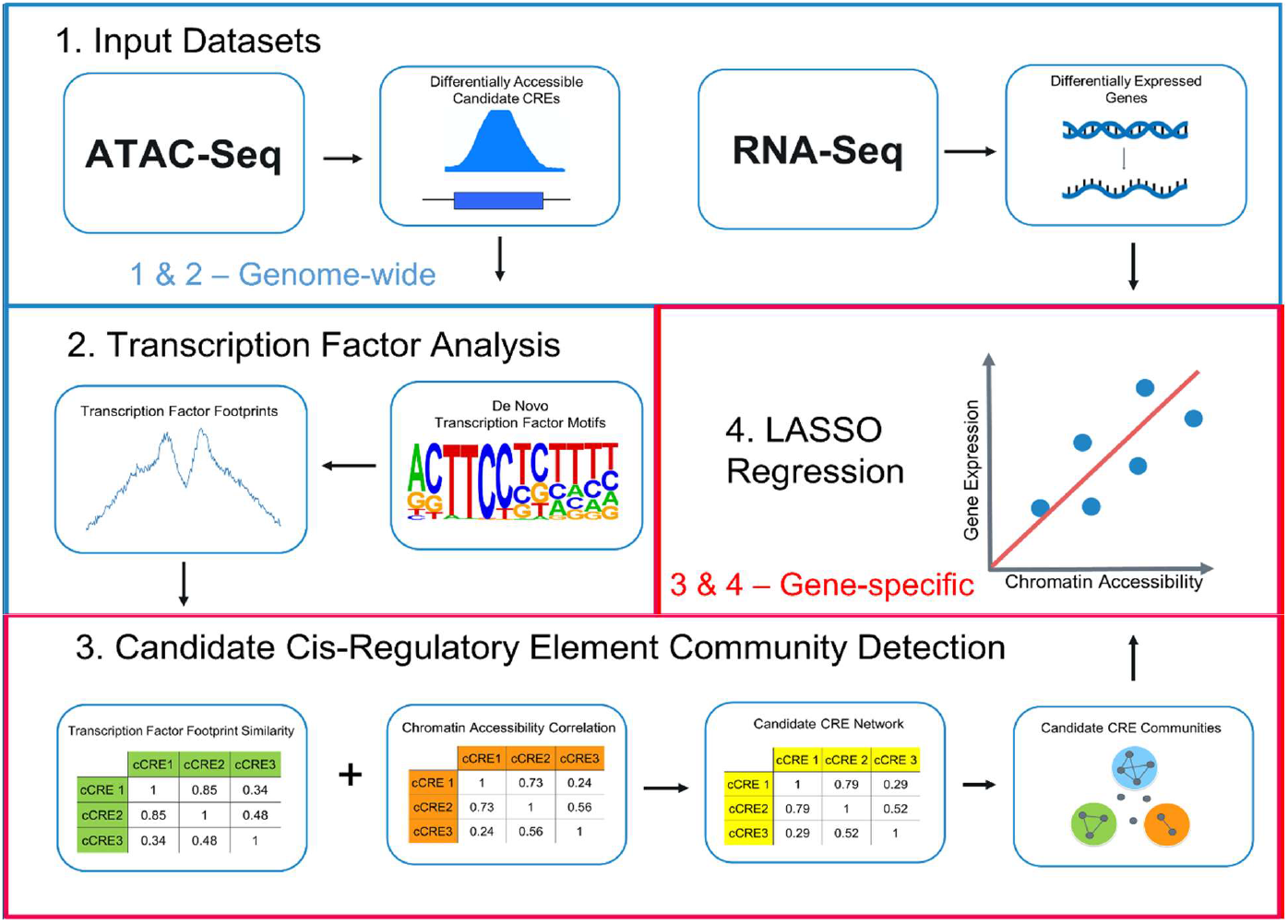
Overview of cisREAD methodology to identify gene-specific cis-regulatory element communities across cellular differentiation from ATAC-seq and RNA-seq datasets. Step 1: candidate CREs are first identified as differentially accessible ATAC-seq peaks and differentially expressed genes are identified for gene-specific modelling. Step 2: enriched transcription factor motifs are curated through *de novo* motif discovery across all genome-wide candidate CREs and matched to transcription factor footprints called in the candidate CREs which are accessible in each differentiation-stage. Step 3: The sample-specific chromatin accessibility of each candidate CRE is characterized, alongside its differentiation-stage specific transcription factor occupancy events and all candidate CREs within 100kb of a target genes TSS, with 3 or more occupancy events, are considered gene-specific candidates. The chromatin correlation and transcription factor similarity of each possible candidate CRE pair is calculated and multiplied to produce integrated similarity scores, used to construct a candidate CRE network. Candidate CREs with similar chromatin accessibilities and transcription factor occupancy events are then grouped together through infomap community detection. Step 4: the chromatin accessibility of each candidate CRE (or mean accessibility of each coCRE) is considered to predict gene expression across all samples in gene-specific LASSO models. LASSO models select candidate (co)CREs whose chromatin accessibility best predicts gene expression and rejects any others. LASSO models are constructed for all differentially expressed genes, and significant predictors of gene expression are considered high-confidence gene-specific predicted CREs.

### Step 1: Inputs

The initial step in the cisREAD workflow is to derive inputs from ATAC-seq and RNA-seq datasets, by finding candidate regulatory elements and protein-coding genes whose activity changes across differentiation. To obtain a set of lineage-specifying candidate CREs (cCREs), we identified differentially accessible regions (DARs) across the time-course using a likelihood ratio test in DESeq2 – comparing a model with Cell-Stage + Donor to a model with Donor only – and retained 97,707 ATAC-seq peaks with a Benjamini-Hochberg (BH) adjusted p-value < 0.01 (Table S1). Similarly, we identified 9,082 differentially expressed genes (DEGs), annotated as protein coding by Ensembl, using a likelihood ratio test, selecting those with a BH adjusted p-value < 0.001 (Table S2).

### Step 2: Transcription factor binding site analysis

cisREAD focuses on identifying gene-specific CREs, targeted by a core set of lineagespecific transcription factors. To discover these TFs, we employed a bottom-up approach using *de novo* motif discovery. HOMER findMotifsGenome.pl was used to find non-redundant DNA k-mers which were over-represented in differentially accessible regions (n = 97,707) compared to constitutively accessible ATAC-seq peaks (n = 146,654). 13 *De novo* motifs, representing unique TF binding sites, were considered enriched by the criteria of statistical over-representation (p < 0.01), frequent occurrence (present in > 2.5% cCREs) and non-redundancy. *De novo* motifs were then matched by HOMER to known TF position-weight matrices (PWMs) from JASPAR Vertebrate Core 2020 (23), HOCOMOCO v11 (24), and HOMER itself (25).

We next identified binding sites for enriched TFs using ATAC-seq footprinting, looking for dips in the ATAC-seq signal where Tn5 transposase cleavage is blocked by TF occupancy (26). TF footprinting was performed with HINT-ATAC, a computational footprinting tool tailored towards ATAC-seq data, to derive differentiation-stage specific TF occupancy events. As input to HINT-ATAC we prepared a differentiation-stage specific BAM file (by merging the BAMs of biological replicate samples with Samtools) and a differentiation-stage specific peak file (by intersecting differentially accessible regions with consensus ATAC-seq peaks accessible in each stage with bedTools). The resultant differentiation-stage specific footprints were then scanned with PWMs of enriched *de novo* motifs, using HOMER findMotifsGenome.pl in -find mode, to predict TF occupancy at each stage. Finally, we summarized the predicted TF occupancy events for genome-wide cCREs in a binary *m* x *n* matrix using a custom python script, where cCREs (n) are scored 1 or 0 to denote the presence or absence of a TF footprint (m). Transcription factor footprints were also visualized as line-plots, showing the mean ATAC-seq signal (adjusted for Tn5 cutting-bias) in the 200bp window centred on each occupied motif in day 0, day 3, day 6 and day 13.

Recognizing that the identification of footprints can be difficult for some transcription factor families, we also employed a footprint-independent method (BMO) for comparison with HINT-ATAC. Here TF occupancy was predicted based on the principle that transcription factors bind motifs that are both frequently occurring and highly accessible (27). BMO was supplied with genome-wide motif scans for each *de novo motif* (obtained using HOMER scanMotifGenomeWide.pl), differentiation-stage specific BAM files and differentiation-stage specific consensus peaks to predict TF occupancy in each differentiation stage.

### Step 3: Candidate CRE network construction and community detection

In steps 1 and 2, we identified DARs across differentiation and characterized their chromatin accessibility and transcription factor occupancy in each cell stage. In steps 3 and 4 we consider each DAR to be a candidate cis-regulatory element, with potential to regulate transcription of one or more differentially expressed genes, also identified in step 1.

To match TF-bound candidate CREs to potential targets, we retained DARs within 100,000 base pairs of a DEG’s transcription start site (TSS), obtained from the Ensembl Canonical Transcript collection (Ensembl Release 104), and discounted those with fewer than 3 TF occupancy events across all time-points. A total of 35,466 DARs were considered as candidate regulators of 8,514 DEGs.

For each gene, we quantified the TF occupancy similarity (Dice coefficient) and chromatin accessibility correlation (Pearson coefficient) between every pair of *n* candidate CREs matched to the gene, and multiplied these measures to produce an integrated similarity score, represented in an *n* x *n* matrix. From this integrated similarity matrix, an undirected weighted graph was produced. This graph represents a gene-specific candidate cis-regulatory network, where nodes are candidate CREs, and edges are candidate CRE pairs, weighted by their integrated similarity score. Edges were only drawn between candidate CREs with integrated similarity scores > 0.3.

From the candidate CRE network, we performed community detection to identify tight-knit groups of densely connected nodes (candidate CREs), which are loosely connected to other nodes in the network. This step groups together candidate CREs which are occupied by similar TFs and accessible in similar differentiation stages, suggesting a co-regulatory mode of gene regulation. Community detection also serves to reduce multicollinearity prior to LASSO regression (step 4), when several candidate elements have correlated chromatin accessibility.

To find communities of co-regulating candidate CREs (coCREs), we implemented Infomap community detection in igraph. Infomap community detection is an optimal, information theory approach using random walks, which performs well on small networks with few intercommunity connections (28),(29).

For each DEG, community detection grouped a mean of 8.4 candidate CREs (range 2-34) into a total of 6.9 (range 1-26) coCREs and lone CREs. coCREs comprised a mean of 2.7 candidate CREs (range 2-12).

### Step 4: Selection of cis-regulatory elements through LASSO linear regression

Following community detection, LASSO linear regression models were constructed for each gene, where the chromatin accessibility of a candidate CRE (or the averaged chromatin accessibility of a coCRE) predicts gene expression across all samples. LASSO is used for variable selection; (co)CREs which do not predict gene expression (i.e. whose chromatin accessibility is not correlated with gene expression) are assigned coefficients of zero and eliminated from the model. LASSO’s capacity for variable selection narrows down the list of candidate (co)CREs to identify those whose cell-specific accessibility best predicts transcription.

As inputs to LASSO regression, we constructed a predictor matrix of independent variables, giving the standardized (z-score) log2 transformed normalized ATAC signal for each (co)CRE within 100kb in all 19 samples, and a response vector of the independent variable, giving the standardized log2 transformed normalized expression of the modelled gene in all samples.

To select the candidate (co)CREs in the predictor matrix which best predict gene expression, we performed LASSO regression for variable selection using the glmnet R package (30, 31). Here a penalty term, equal to the absolute magnitude of their coefficient (*β*) is applied to the regression coefficients, enabling shrinkage to zero and elimination of non-predictive variables from the model. The degree of shrinkage is determined by the regularization parameter *λ*. We determined the optimum value of *λ*, which minimizes the minimum mean cross-validated error of the model (*λ_min_*), through 10-fold random cross-validation (Figure S2) and then constructed the final LASSO model at *λ_min_* to perform variable selection. All candidate CREs with regression coefficients equal to zero were considered rejected, and all remaining variables with ≠ 0 were considered selected. Selected variables, with *β* ≠ 0, were then subject to significance testing using the selectiveInference R package using a method which adjusts for how predictors were cherry-picked from a larger pool of candidates (32).

To account for multiple testing and control the False Discovery Rate (FDR), gene-specific p-values were assigned to each model (equal to that of its most significant predictor) and subject to BH-adjustment (33). Candidate (co)CREs with p-values < 0.05, and gene-specific adjusted p-values < 0.05, were considered statistically significant predictors of gene expression.

A total of 24,648 candidate CREs were selected by at least one model to regulate a DEG, with a mean of 4.7 candidates (range 1 to 22) selected per gene. CREs were selected for 8,215 (96%) of modelled genes, with the mean CRE selected for 1.6 genes (range 1-12). A total of 20,818 CREs were rejected by all models in which they were potential predictors and are therefore determined not to regulate a differentially expressed gene.

24,181/38,544 (63%) selected CRE-gene relationships were characterized by positive correlation of chromatin accessibility with gene expression, indicating upregulation, and 13,368 (37%) were characterized with negative correlation, indicating downregulation. 6,063/9,440 (64%) of significant CRE-gene relationships (64%) indicated upregulation, and 3,377 (36%) indicated downregulation.

8,022 selected CREs (33% of selected, 24% of candidates), were found to be statistically significant predictors of gene expression in at least one model, following correction for multiple testing. 4,166 (51% of models) were found to have at least one statistically significant predictor, with an average of 2.6 significant CREs per model (range 1-16). Each significant CRE was a statistically significant predictor in a mean of 1.2 models (range 1-6). We considered cisREAD-significant CREs to represent a top-tier set of gene-specific regulatory element predictions, prioritized across B cell differentiation.

### Downstream analysis of cisREAD results

To gain biological insight into predicted gene regulation, we employed a number of additional bioinformatics analyses. These included clustering chromatin accessibility of cisREAD-predicted CREs, and relating target genes to Parsimonious Gene Correlation Network Analysis (PCGNA) co-expression modules (34). To check that conclusions are robust to binding site prediction method, we also ran cisREAD replacing footprinting in step 2 with the footprint-independent BMO method (27) and repeated downstream analyses. Predicted occupancy was also compared with ChIP-seq data for PU.1/SPIB and AP-1 factors. Full details are provided in Supplementary Methods and Figures.

## RESULTS

### A core transcription factor network occupies differentially accessible chromatin regions

We identified 97,707 differentially accessible regions (DARs) as genome-wide candidate CREs (Table S1). Transcription factor binding site analysis revealed the enrichment of 13 *de novo* motifs in DARs, which were mapped to transcription factor footprints at 9 differentiation stages (Figure 3). The TFs grouped under each motif, represent a core set of transcription factors that drive the differentiation of mature B cells to plasma cells.

**Figure 3.**
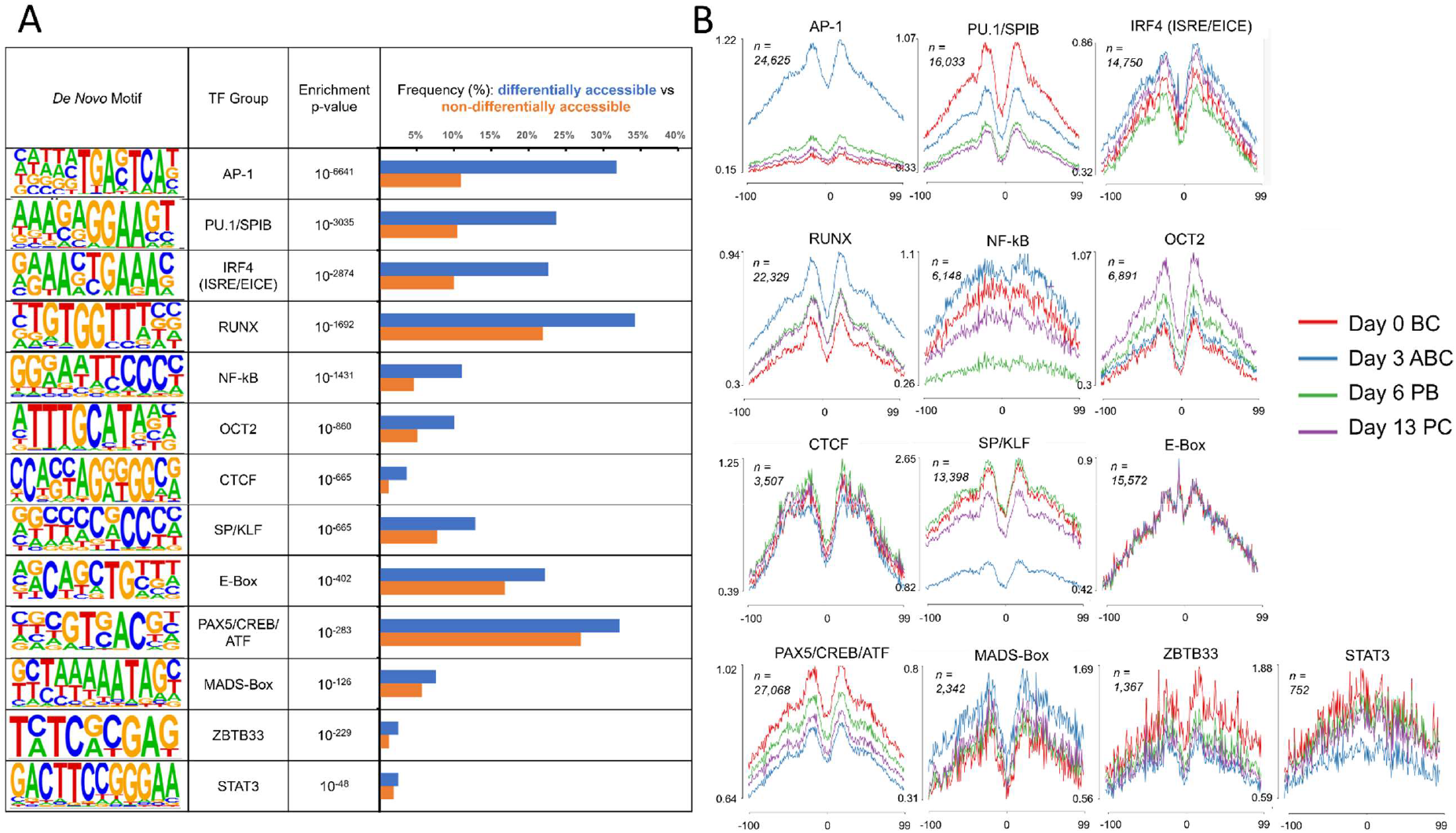
A) Enriched de novo motifs discovered using HOMER in genome-wide candidate CREs (differentially accessible ATAC-seq peaks) compared to universal, non-differentially accessible ATAC-seq peaks. Statistically enriched, frequently occurring and non-redundant de novo motifs (represented by sequence logos) are curated into a set of ‘core transcription factors’ which represents the dominant transcription factors responsible for gene regulation across plasma cell differentiation. B) ATAC-seq footprints called in each differentiation stage by HINT-ATAC, matched to enriched de novo motifs. Line plots show the ATAC-seq signal (y-axis) for the 200 base-pair window centred on the transcription factor footprint (x-axis), averaged across all footprints called in a differentiation stage. Footprints are shown for day 0 B cells, day 3 activated B cells, day 6 plasmablasts and day 13 plasma cells. N gives the total number of transcription factor footprints called over the four stages.

The most significantly enriched *de novo* motif matched the binding site for the AP-1 transcription factor complex. The mean ATAC-seq signal surrounding AP-1 footprints (indicating the number of ATAC-seq fragments which mapped to the DAR) was high in day 3 ABCs, and low in BCs, PBs and PCs, indicating that accessibility was linked to the gain and loss of CD40 and BCR activating signals. PU.1/SPIB ETS factor motifs were also highly significantly enriched in DARs.

PU.1/SPIB footprint accessibility declined upon B cell activation, and then declined further following the plasmablast transition. We also observed enrichment of a de novo motif matching IRF4 binding sites at interferon-specific response elements (ISREs) and ETS-IRF composite elements (EICEs). Unlike AP-1 and PU.1/SPIB, IRF4 (ISRE/EICE) footprints were highly accessible throughout the time-course, yet accessibility increased following activation, decreased upon the PB transition, and increased again in PCs.

RUNX motifs occurred most frequently out of all de novo motifs, and also gained accessibility following B cell activation, which declined slightly in plasmablasts and plasma cells. NF-κB motifs were also enriched in DARs and exhibited a ‘shallow’ footprint which gained accessibility following CD40 and BCR engagement, lost accessibility following loss of CD40L, and regained accessibility following addition of APRIL at day 6. OCT2 footprint accessibility did not change upon B cell activation, however accessibility was gained following the plasmablast transition and further increased upon APRIL-driven PC differentiation.

CTCF motifs were enriched in DARs yet were infrequent, occupying just 3% of CREs. CTCF displayed a ‘deep’ footprint, which was similarly accessible across all differentiation stages. SP/KLF footprints exhibited high accessibility in all cell states apart from ABCs; accessibility was lost upon B cell activation and regained following removal of activation stimuli. E-Box footprints were similarly accessible in all cell stages. A de novo motif matching binding sites for PAX5, CREB and ATF factors was also enriched, which gained accessibility alongside B cell activation, and lost accessibility alongside plasma cell differentiation. MADS-Box footprints were most accessible in activated B cells. ZBTB33 and STAT3 motifs were enriched in DARs yet are infrequent, resulting in noisy footprint profiles where accessibility changes were unclear. The de novo STAT3 motif includes a GAC sequence preceding the consensus TTCCNGGAA motif, and so would match a subset of STAT3 binding site

### Gene-specific models prioritise cis-regulatory elements linked to the expression of differentially expressed genes

In order to link dynamic TF occupancy (Figure 3) to differential gene expression we employed cisREAD to assign differentially accessible regions to target genes. 35,466 transcription factor-bound DARs were considered candidate regulators of differential expression based on their proximity (<100kb) to a differentially expressed gene (Table S3). 24,648 candidate cis-regulatory elements were selected by cisREAD to regulate one or more DEGs, based on correlation between accessibility and expression (Table S4). Of these, 8,022 exhibited temporal accessibility that significantly predicted differential gene expression, with a total of 9,440 CRE-DEG pairs (Table S5). These ‘cisREAD-significant’ CREs are considered to be a top tier set of gene-specific regulatory elements whose dynamic activity is important for B cell differentiation.

cisREAD linked both positive (enhancer) and negative (silencer) regulators to target gene expression, identified by the sign of accessibility-expression correlation. A range of distal and proximal CREs were linked to gene expression, acting across a median distance of 44kb for significant CREs. 44% of CREs significantly linked to gene expression were intronic, whilst 24% were intergenic and 23% were annotated as promoters. A small proportion of significant CREs overlapped exons (6%), 3’ untranslated regions (UTRs) (2%) and 5’ UTRs (1%).

### Cluster analysis of gene-specific cis-regulatory elements reveals TF binding dynamics across differentiation

To interrogate regulatory dynamics on a genome-wide scale, we divided significant genespecific CREs into 8 regulatory clusters based on differentiation-stage specific chromatin accessibility using k-means clustering (Figure 4A). *k* was incremented from 3 onwards until early-stage and late-stage ABC-specific clusters were separated. For each cluster, we then calculated enrichment of core TF footprints (Figure 4B) and de novo motifs (Figure 4C) to characterize the transcriptional regulators which drive gene regulation at each stage.

**Figure 4.**
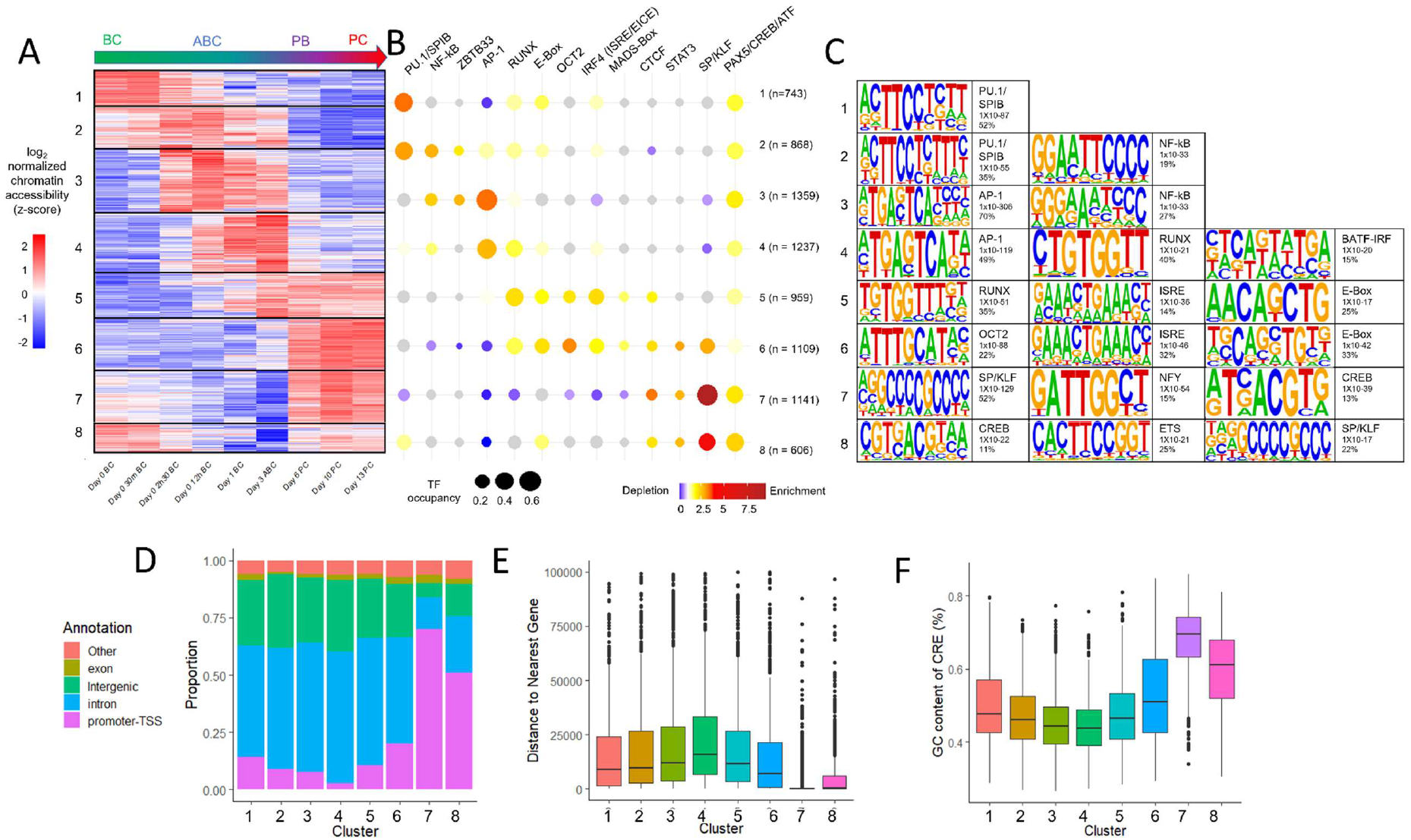
Enrichment of footprints and de novo motifs in cis-regulatory clusters. A) Heatmap showing log2 normalized chromatin accessibility (z-score) of cis-regulatory elements significantly linked to gene expression, k-means clustered (k = 8) by z-score chromatin accessibility. B) Bubbleplot showing enrichment of TF occupancy in each cluster. Size of bubbles gives the proportion of each cluster harbouring a TF footprint, colour shows significant (p < 0.05, two-sided Fisher test) enrichment (fold-change between cluster and other clusters > 1, red) or depletion (fold-change between cluster and other clusters < 1, blue), grey represents no significant enrichment. N gives the number of CREs in each cluster. C) Top 3 de *novo* motifs enriched in each cluster from HOMER. D) Genomic annotation of each CRE by HOMER. E) Boxplot showing distance of CREs to their nearest gene in each cluster. F) Boxplot showing GC content of CREs in each cluster.

Cluster 1 CREs were accessible prior to B cell activation, and lost accessibility at 2h:30-24 hours post activation. 48% of CREs in this cluster harboured a PU.1/SPIB footprint. Cluster 2 CREs were similarly accessible in B cells, yet sustained accessibility after stimulation, remaining accessible until the plasmablast transition at day 6. These B cell specific CREs were similarly enriched in PU.1/SPIB footprints, present in 42% of elements. NF-κB footprints were also enriched in cluster 2, but not cluster 1, indicating association with B cell activation.

CREs which gained accessibility in activated B cells 2h30 post activation were placed in cluster 3. This early-ABC cluster was also associated with occupancy by the AP-1 complex (64% of CREs) and NF-κB. ABC-specific CREs which gained accessibility at 12h post activation were assigned to cluster 4. This cluster was similarly enriched in AP-1 footprints (52% of CREs), also showed enrichment of RUNX footprints (33% of CREs). *De novo* motif discovery found enrichment of a BATF(AP-1)-IRF4 composite motif, not enriched in the earlier cluster 3.

Cluster 5 CREs gained accessibility around 1-3 days after activation, and sustained accessibility throughout plasma cell differentiation. CREs in this cluster were enriched in occupancy by RUNX, E-Box, OCT2 and ISRE binding TFs. *De novo* motif analysis also reported significant enrichment of these factors. Cluster 6 CREs exhibited PB/PC-specific accessibility, and were likewise enriched in occupancy by RUNX, E-Box, OCT2 and ISRE, alongside SP/KLF.

Cluster 7 CREs were also accessible in plasmablasts and plasma cells, yet exhibited moderate accessibility prior to activation. Notably CREs in this cluster were over-represented for gene promoters and highly GC-rich, they exhibited strong enrichment for SP/KLF factors which occupied 66% of the cluster. *De novo* motif analysis revealed enrichment of additional promoter-associated motifs including NFY and CREB which conform to elements of previously defined XBP1 binding motifs (35, 36).

CREs in cluster 8 were accessible in initial B cell populations and plasmablasts and plasma cells, but declined in accessibility during ABC cell stages. Similar to cluster 7, this class was overrepresented for promoters and GC-rich elements, although to a lesser extent (Figure 4D and Figure 4E). SP/KLF footprints were also enriched in this cluster, and *de novo* motif enrichment showed enrichment of CREB/XBP1 and ELK/ETV/ETS factors.

Due to infrequent motif occurrence, we failed to reliably associate STAT3, ZBTB33, CTCF and MADS-Box footprints with any cell-specific regulatory clusters. Furthermore, the broad spectrum of TF binding sites matched by the PAX5/CREB/ATF motifs resulted in enrichment in multiple distinct regulatory clusters, representing motif occupancy by a wide range of differentially active TFs.

In order to alleviate concerns over the reliability of footprinting for transient binding factors we also employed cisREAD using TF binding events predicted by a footprint independent method BMO (described in supplementary methods) (27). The analysis shown in Figure 4 was repeated using cisREAD predictions derived from BMO binding sites and showed highly concordant results (Figure S3).

### Parsimonious Gene Co-expression Network Analysis shows differential transcriptional regulation of temporally distinct biological pathways

To investigate how differential transcription factor occupancy controls gene expression dynamics, we linked cisREAD-identified gene-specific CREs to co-expression modules identified using the Parsimonious Gene Co-Expression Network Analysis (PGCNA) method (34).

PGCNA was performed on the 19 RNA-seq samples from Figure 1, alongside 3 additional samples (day 0 6h x 2, day 6 x1) without matching ATAC-seq samples. A total of 23 gene co-expression modules were identified, each enriched in differential biological processes. For the 10 largest modules (M1-10), we identified cis-elements that significantly predicted gene expression in each module, and calculated enrichment of the 13 transcription factor footprints. Interactive gene co-expression networks can be accessed at https://matthewcare.wixsite.com/pgcna/bcell-detailedtc.

To look specifically at transcriptional activation by each TF, we identified significant coCRE predictors in each model that are positively correlated with gene expression. We calculated enrichment of transcription factor footprints in cis-elements (Figure 5A) associated with coexpression modules (Figure 5B), identifying regulatory inputs into basal gene expression across differentiation.

**Figure 5.**
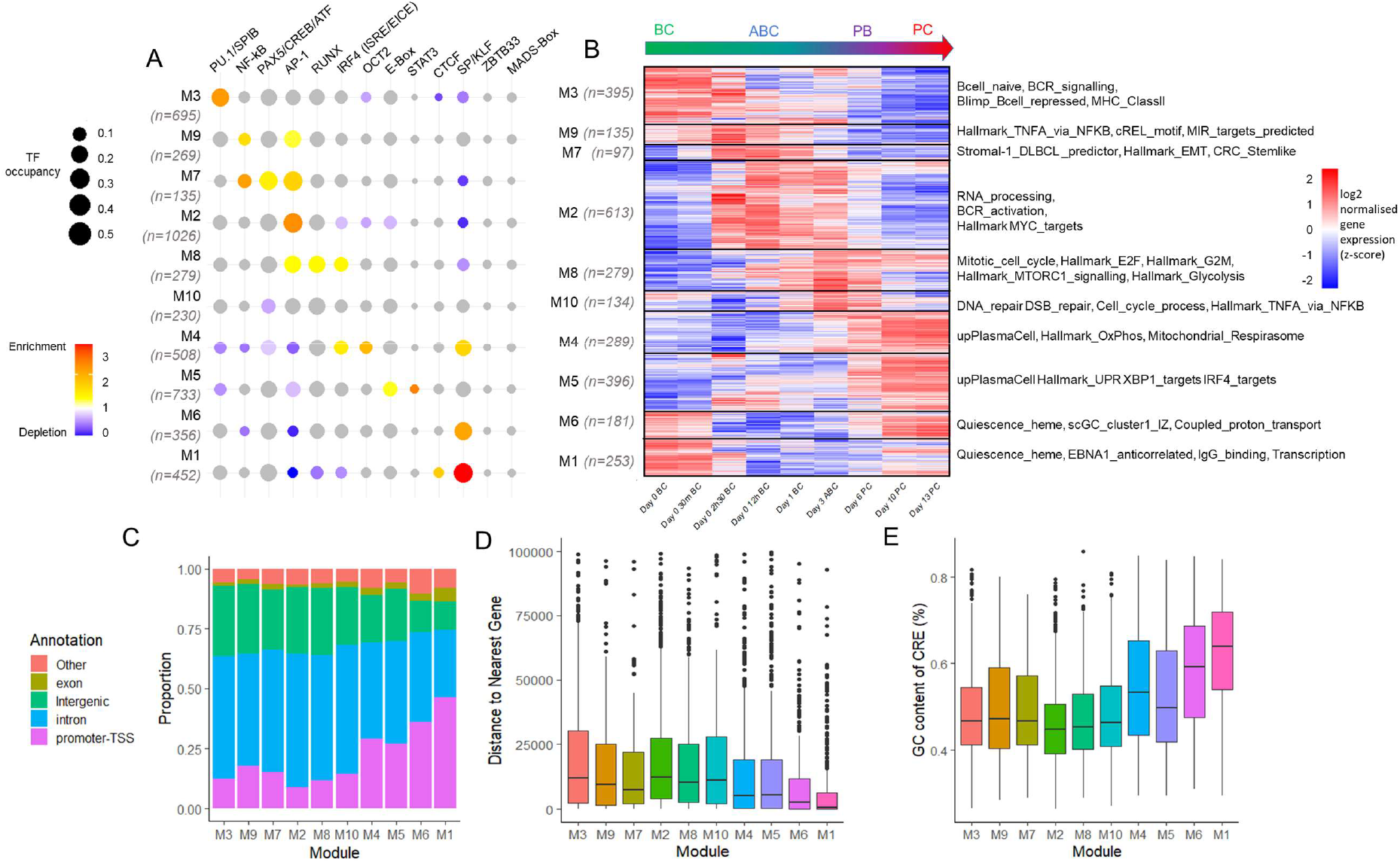
Enrichment of footprints in PGCNA co-expression modules. A) Bubbleplot showing enrichment of TF occupancy in each cluster. Size of bubbles gives the proportion of each cluster harbouring a TF footprint, colour shows significant (p < 0.05, two-sided Fisher test) enrichment (fold-change between genes in module and genes not in module > 1, red) or depletion (fold-change < 1, blue), grey represents no significant enrichment. N gives the number of genes with significant CREs in each module. B) Heatmap showing log2 normalized gene expression (z-score) of genes with significantly linked CREs, module names reflect enriched gene sets in each module. C) Genomic annotation of each CRE by HOMER. D) Boxplot showing distance of CREs to their nearest gene in each cluster. E) Boxplot showing GC content of CREs in each cluster.

We observed an association between PU.1/SPIB occupancy and B cell-specific gene expression in module 3. This module contained 1,523 genes enriched in gene sets relating to naïve B cells, BCR signalling, repression by BLIMP1 and the major histocompatibility (MHC) Class II complex. 695 cis-regulatory elements were linked to the expression of 395 or 25% of genes in this module, whose expression declined following B cell activation. 38% of CREs regulating this B cell specific module featured PU.1/SPIB footprints, over twice the proportion of PU.1/SPIB footprints in other modules. Importantly, the enrichment of PU.1/SPIB in B cell specific gene expression modules, mirrored the enrichment of PU.1/SPIB in B cell specific chromatin accessibility clusters shown in Figure 4.

NF-κB footprints were significantly enriched in CREs linked to module 9, where genes were expressed in both B cell and activated B cell stages, peaking 2h30 post-activation. Module 9 genes (n = 768) were specifically associated with regulation by NF-κB, overlapping validated NF-κB target genes (Hallmark_TNFA_via_NFKB) and genes with nearby NF-κB motifs (cREL_motif). cisREAD linked 135 (17%) genes in this module to activation by 269 regulatory elements, 12.6% of which harbored an NF-κB footprint. AP-1 footprints were also significantly enriched in this module, suggesting that both AP-1 and NF-κB regulate expression in the initial stages of B cell activation.

AP-1 enrichment was observed across modules 7, 2 and 8, indicating sustained regulatory input into sequential ABC-specific transcriptional programs. This reflects observed AP-1 occupancy at early, mid and late chromatin accessibility clusters in Figure 4. Alongside sustained AP-1, differential TF co-enrichment was observed in early and late ABC coexpression modules.

NF-κB enrichment continued from module 9 into module 7, where 969 genes were upregulated in the earlier stages of B cell activation. PAX5/CREB/ATF footprints were also enriched in this module, which was associated with Stromal-1_DLBCL _predictor, Hallmark_EMT and CRC_Stemlike gene-sets. In contrast, AP-1 was the only transcription factor associated with the following ABC-specific module 2, hosting 1,749 genes enriched in RNA processing, BCR activation and regulation by MYC. Alongside AP-1, RUNX and IRF4 (ISRE/EICE) footprints were enriched in CREs linked to genes in module 8 (n = 817), expressed in both ABCs and PBs. These genes were enriched in terms relating to the cell cycle, MTORC1 signalling, glycolysis and regulation by E2F. RUNX and IRF4 were previously observed to occupy CREs accessible in both ABCs and PBs/PCs. Whilst no TF footprints were enriched in DNA-repair-associated ABC/PB module 10, this was also the smallest module.

Modules 4 and 5 both included genes upregulated in plasmablasts and plasma cells. IRF4, OCT2 and SP/KLF footprints were enriched in CREs linked to module 4 genes (n = 1,293), associated with oxidative phosphorylation and mitochondrial respirasome pathways. However, module 5 genes (n = 1,227), associated with XBP1 and IRF4 targets and the unfolded protein response, were instead linked to CREs enriched in E-Box and STAT3 footprints. Whilst the analysis in Figure 4 linked occupancy of IRF4, OCT2, SP/KLF, E-Box and STAT3 factors to accessibility in PBs/PCs, differential module enrichment would suggest divergent regulatory functions in terminal plasma cell differentiation.

Modules 6 (n = 1,174) and 1 (n =2,154) host genes which were upregulated in non-dividing B cells and plasma cells and were both enriched for heme_quiescence gene sets. CREs linked to both modules were significantly enriched for SP/KLF footprints, which were found in 37.4% of module 6 CREs and 47.6% of module 1 CREs. Notably, CREs linked to these modules included more promoters (Figure 5C), more gene-proximal regions (Figure 5D) and more GC-rich sequences (Figure 5E), compared to CREs linked to other modules.

In summary our results show that PU.1/SPIB is associated with B cell specific expression, NF-kB and AP-1 link to modules associated with activated B cell expression, RUNX, IRF4, OCT2, E-Box and STAT3 are linked to plasmablast and plasma cell expression and SP/KLF associates with quiescent B cell and plasma cell expression. A repeat of the analysis using BMO-derived cisREAD predictions yielded similar associations (Figure S4).

### Gene regulation shifts from PU.1/SPIB to AP-1 during B cell activation

In Figures 3, 4 and 5 we observed an association between PU.1/SPIB with the B cell state, and AP-1 with the activated B cell state. To further investigate how the shift in regulatory inputs shapes B lineage expression programmes, we used cisREAD to predict PU.1/SPIB and AP-1 target genes and examine how transcription factor occupancy drives gene expression at distinct stages of differentiation.

We first identified all genes predicted by cisREAD to be upregulated by CREs with PU.1/SPIB or AP-1 footprints (Figure 6C). Predicted PU.1/SPIB and AP-1 target genes were collectively clustered by their expression using k-means clustering (k=5, incremented until early-stage and late-stage ABC expression clusters were separated). Target genes were annotated by predicted regulation; by linkage to CREs with PU.1/SPIB footprints (orange), AP-1 footprints (pink) or both PU.1/SPIB and AP-1 footprints (either in the same CRE or in separate CREs, blue). Alongside these genes, we observed the chromatin accessibility of their cisREAD-predicted CREs (Figure 6B). Similarly, cis-elements were annotated by the presence of footprints for PU.1/SPIB only (orange), AP-1 only (pink), and both PU.1/SPIB and AP-1 at the same CRE (blue).

**Figure 6.**
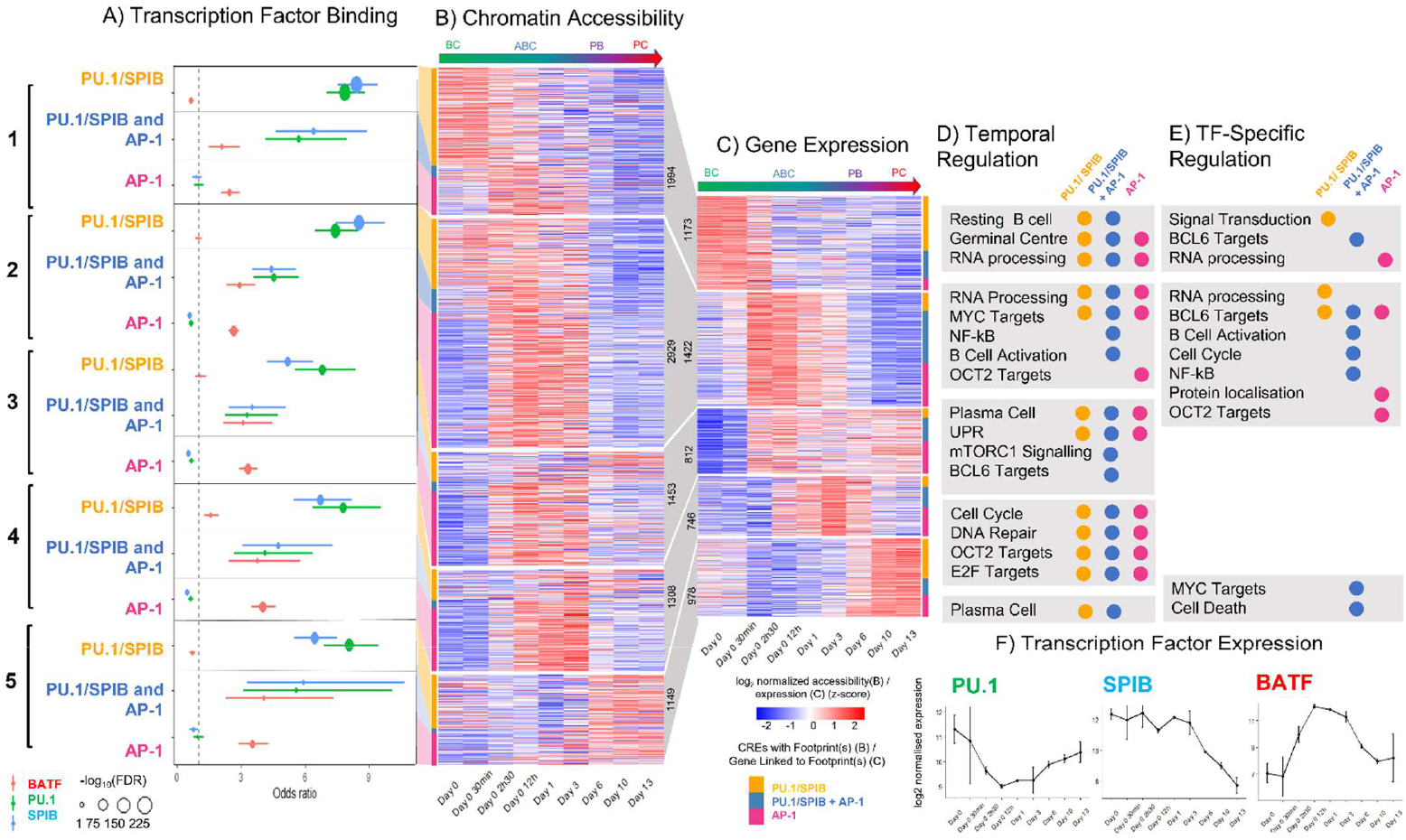
PU.1/SPIB and AP-1 (BATF) occupancy at footprinted cis-regulatory elements linked to gene expression clusters. Starting at C) Heatmap showing the log2 normalized expression of genes linked y cisREAD to CREs with PU.1/SPIB and/or AP-1 footprints, k-means clustered by expression (k=5). Genes are annotated based on predicted regulation by PU.1/SPIB only (yellow), PU.1/SPIB and AP-1 (blue, either in the same CRE or at different CREs) or AP-1 only (pink). B) Heatmap showing the log2 normalized chromatin accessibility of CREs linked to gene expression clusters, annotated by the presence of footprints for PU.1/SPIB only (yellow), PU.1/SPIB and AP-1 at the same CRE (blue) or AP-1 only (pink). A) Enrichment of ChIP-seq binding sites for SPIB (from OCILY-3/OCILY-10, blue), PU.1 (from GM12878, green), and BATF (an AP-1 factor, from GM12878, red) at CREs with footprints for PU.1/SPIB and/or AP-1, linked to each expression cluster. Enrichment is calculated by two-sided fisher test, comparing ChIP-seq binding for each factor at CREs, in each cluster, with PU.1/SPIB footprints only (yellow), PU.1/SPIB and AP-1 footprints (blue) and AP-1 footprints (pink) to all DARs without the given footprint. D) Gene sets enriched (one-sided fisher test, FDR < 0.1) in genes in each cluster, linked to regulatory elements with PU.1/SPIB (orange), PU.1/SPIB + AP-1 (blue) and AP-1 (red) footprints, compared to genes in other clusters. E) Gene sets enriched (one-sided fisher test, FDR < 0.1) in genes in each cluster, linked to regulatory elements with PU.1/SPIB (orange), PU.1/SPIB + AP-1 (blue) and AP-1 (red) footprints, compared to genes with similar expression patterns, not linked to given footprints F) log2 normalized PU.1, SPIB and BATF expression.

We initially assessed each cluster in Figure 6C by the proportion of genes predicted to be upregulated by PU.1/SPIB or AP-1 uniquely or in combination. We observed that most B cell upregulated genes were linked to PU.1/SPIB footprints, which were most accessible in B cell time-points (Figures 6B and 6C cluster 2). Conversely, most ABC upregulated genes were linked to AP-1 footprints which were accessible at ABC time-points (Figures 6B and 6C clusters 3 and 4). Almost half of genes upregulated in the hours following CD40 and BCR stimulation were predicted targets for both PU.1/SPIB and AP-1 (Figure 6C cluster 2). PU.1/SPIB and AP-1 were predicted to regulate expression of these genes at different sets of regulatory elements, as indicated by the comparatively small set of cis-elements with both PU.1/SPIB and AP-1 footprints (cluster 2 blue bar in Figures 6B and 6C).

To confirm transcription factor occupancy at PU.1/SPIB footprints we employed existing PU.1 ChIP-seq data from lymphoblastoid cell line GM12878 (37), and SPIB ChIP-seq data from Diffuse Large B Cell Lymphoma (DLBCL) cell lines OCILY-3 and OCILY-10, which are surrogates for different activated B cell states (38). Both PU.1 and SPIB ChIP-seq binding sites significantly overlapped CREs with PU.1/SPIB footprints, but not CREs with only AP-1 footprints (Figure 6A). Furthermore the effect size of PU.1 and SPIB enrichment varied by cluster; PU.1 binding sites were most enriched at CREs with plasmablast/plasma cell accessibility (linked to clusters 3, 4 and 5) and SPIB binding sites were most enriched at CREs with activated B cell accessibility (linked to clusters 1 and 2). This may reflect increased *SPIB* in activated B cells and increased *PU.1* mRNA in PBs/PCs (Figure 6F).

Amongst potential AP-1 binding transcription factors BATF has been previously identified as a regulator of germinal centre gene expression and a key driver of neoplastic B-cells related to the activated B-cell state, equivalent to day 3 of *in vitro* differentiation (41). To test BATF enrichment at AP-1 footprints we employed existing BATF ChIP-seq data from GM12878 (37) and found significant enrichment of BATF binding sites at CREs with AP-1 footprints, but not at CREs with only PU.1/SPIB footprints. Mirroring transcription of *BATF,* we observe greatest enrichment of BATF binding sites at CREs most accessible 12 hours to 3 days after activation (linked to cluster 4).

To suggest functions for PU.1, SPIB and AP-1 (BATF) in the B cell to activated B cell transition we performed gene set over-representation analyses, identifying gene sets enriched in transcription factor target genes (PU.1/SPIB only, PU.1/SPIB and AP-1, and AP-1 only) in each cluster. Firstly, to identify pathways enriched by differentiation stage we first used a background of genes targeted by any of the three factors in any other cluster. ‘Temporally enriched’ gene sets (FDR < 0.1) are summarized on Figure 6D and listed in full in Table S6. Secondly, to identify pathways preferentially regulated by PU.1/SPIB and/or AP-1 at each stage we used a background of similarly expressed genes, not linked to the given footprint(s) (Figure S5). These ‘TF-specific enriched’ gene sets are summarized on Figure 6E and listed in full in Table S7.

For cluster 1, PU.1/SPIB, but not AP-1, targets were enriched for B cell specific gene sets (D), showing TF-specific enrichment of genes involved in intracellular signal transduction (E). Both PU.1/SPIB and AP-1 were enriched for germinal center expressed and RNA processing gene sets (D) and AP-1 targets were enriched for RNA metabolism genes, compared to similar genes not linked to AP-1 (E).

A variety of gene sets relating to RNA processing and MYC targets (PU.1/SPIB and AP-1), NF-κB and BCR activation (PU.1/SPIB in conjunction with AP-1) and OCT2 targets (AP-1 only) were enriched in cluster 2, over all over clusters (D). RNA-processing genes showed preferential regulation by PU.1/SPIB, and OCT2 targets showed preferential regulation by AP-1 (E). A number of gene sets showed preferential co-regulation by PU.1/SPIB and AP-1, including NF-κB pathway, cell cycle and BCL6 target genes (also observed for cluster 1) (E).

Genes expressed later in the activation process (cluster 4) were enriched for gene sets relating to cell cycle and DNA repair, alongside OCT2 and E2F targets, regardless of regulator. No transcription factor specific pathways were enriched for this cluster.

Overall this combined view of PU.1/SPIB and AP-1 target genes suggests that gene regulation shifts from PU.1/SPIB to AP-1 during B cell activation, contingent on each factors’ induction, with potential coregulation of activation pathways during the immediate response to CD40 and BCR engagement.

## DISCUSSION

B cell differentiation is driven through changes in genetic regulation, where regulatory circuits are rewired through dynamic shifts in epigenetic remodelling and transcription factor activity. Genome wide studies of epigenomic, transcriptomic and conformational datasets have been instrumental in uncovering how the chromatin environment shapes B cell populations, and responds to stimuli to determine cell fate (5, 8, 10, 35, 39, 40). However, there are gaps in our understanding of the complex associations between *cis* and *trans* acting factors which fine-tune transcription to orchestrate B cell maturation.

Our fine-grained map of chromatin accessibility and gene expression profiles across B cell differentiation has allowed us to identify stimuli-responsive shifts in transcription factor binding associated with distinct epigenetic programmes. Key to these findings has been our new method; a data-driven approach which prioritises dynamically accessible regulatory elements, targeted by lineage-specific transcription factors and associated with the expression of differentiation associated genes. Application of our method to a model system of human B cell differentiation has revealed how a core network of transcription factors exercise regulatory control over B cell differentiation, explicitly coupled to the CD40 and BCR activation stimuli administered to the *in vitro* cell system.

In many cases, observed regulation downstream of CD40 and BCR signals is consistent with regulatory dynamics observed in *ex vivo* murine B cells stimulated by lipopolysaccharide (LPS) (5, 39, 40) as well as in primary human B cell populations (8, 10). Findings reported in these studies, in conjunction with the analyses presented here, suggests that common mechanisms define genetic regulation during B cell activation and plasma cell differentiation; characterised by declining regulation at PU.1/SPIB associated motifs and increased regulation at motifs occupied by NF-κB, AP-1, IRF4, OCT2, E2A (E-Box) and MEF2 factors (MADs-Box). Throughout the results presented here, we repeatedly see that the B cell to activated B cell transition is defined by a shift from PU.1/SPIB-driven to AP-1-driven gene regulation.

PU.1 and SPIB are two partially redundant transcription factors, shown to upregulate BCR signal transduction and receptors for CD40L, BAFF and TLR ligands and are required for B cell activation (41). *PU. 1* and *SPIB* are both downregulated in plasma cells, and SPIB overexpression is associated with germinal center subtype DLBCL (38). PU.1/SPIB footprints were uniquely enriched in B cell specific cis-regulatory clusters (Figure 4) and linked to the expression of B cell specific gene modules expressed at these time-points (Figure 5). PU.1/SPIB enriched modules (Figure 5), and PU.1/SPIB target genes (Figure 6) were enriched in pathways relating to B cell expression, BCR signalling and signal transduction. This is consistent with PU.1 and SPIB’s complementary roles in environment sensing to facilitate B cell activation (41).

Our data suggest that some PU.1/SPIB footprints sustain accessibility after activation (Figures 3, 4 and 6) and may contribute regulation to genes immediately induced by activation stimuli (Figure 6). Unlike *PU. 1,* we found that *SPIB* expression is maintained postactivation (Figure 6F) and that SPIB may preferentially occupy PU.1/SPIB motifs to modulate transcription (Figure 6A).

AP-1 occupied cis-regulatory elements open following CD40 and BCR engagement (Figures 3, 4 and 6). AP-1 provides regulatory input from the onset of activation until the plasmablast transition, and gene set enrichment results suggest diverse functions controlling B cell activation, RNA processing and the cell cycle (Figures 5 and 6).

AP-1 subunits (including FOS, FRA1 or BATF partnered with JUN, JUNB or JUND) regulate temporally diverse processes in B cell maturation (42–48). BATF is induced in a CD40, and MHC-II dependent manner (48, 49), and is essential for co-ordinating class-switch recombination (CSR) and germinal center establishment (46, 50). In germinal centers, BATF is induced as B cells transition from the light zone to dark zone upon selection by T cells and its expression is associated with cell cycle re-entry (48). BATF over-expression is associated with ABC-subtype DLBCL (38).

Our data show that BATF is induced following CD40 engagement, and BATF ChIP-seq binding sites for GM12878 overlap AP-1 footprints accessible in activated B cells (Figure 6). AP-1 functions suggested by gene set over-representation (Figures 5 and 6) support known roles for BATF in class switch recombination (DNA repair) and cell cycle entry, and indicate additional involvement in RNA processing, crosstalk with the NF-κB pathway and regulation of MYC, E2F and OCT2 target genes.

In Figure 6 we see evidence that the transition from PU.1/SPIB-driven to AP-1 driven regulation is graded by expression of *PU. 1, SPIB* and *BATF.* Our data suggest that the two alternate transcriptional programs intersect in the hours following activation, when NF-κB, MYC and BCL6 are induced (51–54). We see significant over-representation of NF-κB, MYC and BCL6 targets in both cisREAD-predicted PU.1/SPIB and AP-1 targets. This suggests that, at this transitory stage, PU.1/SPIB and AP-1 regulatory networks are intertwined with those of other critical germinal center factors. Overall our data support a model where gene regulation gradually shifts from PU.1/SPIB towards AP-1 upon B cell activation, passing through an intermediate stage where SPIB and BATF may co-ordinate the expression of genes induced immediately upon activation.

Altogether we demonstrate that cisREAD successfully prioritises differential transcription factor occupancy and chromatin accessibility with dynamic gene expression across B cell maturation. Applying this method to our *in vitro* system we were able to reveal new insight into transcriptional reprogramming during B cell activation. However, both the dataset and methods have their limitations.

Firstly, due to preferential differentiation of memory B cells into long-lived plasma cells, whilst day 0-3 time-points originate from total peripheral blood B cells, day 6-13 time-points originate from isolated memory B cell subpopulations. Since we focus interpretation primarily on factors associated with the BC-ABC transition, differences between total B cell derived and memory B cell derived gene regulation should not affect these conclusions.

Secondly, in the absence of measured transcription factor binding, we use transcription factor footprinting, corrected for Tn5 cutting bias, as a proxy for TF occupancy at accessible regions (26). Whilst many studies successfully employ ATAC-seq footprinting to interrogate TF binding dynamics (39, 55, 56), it has been noted that many transcription factors do not leave strong footprints, particularly those with short DNA residency times (27, 57, 58). This is particularly a concern for transiently binding signal-inducible TFs like STAT3 and NF-κB, for which we observe detectible yet ‘shallow’ footprints (Figure 3B). The replication of our analysis with binding site predictions derived from the BMO model, based on motif accessibility and co-occurrence (27), suggests however that our cisREAD method and downstream analysis is robust to the use of footprints (Figures S3 and S4).

We also note that we have not been able to differentiate specific motifs or footprints for two key transcriptional regulators of terminal PC differentiation PRDM1 and XBP1. Both of these factors are expressed and occupy target sites in differentiating plasma cells (35). PRDM1/BLIMP1 motifs overlap with a subset of ISREs and may therefore be subsumed amongst a subset of accessible regions with an ISRE match. We identified additional binding motifs consistent with the XBP1 and associated NFY consensus sequences in CRE clusters 7 and 8 (5-ACGTG-3/5-CACGT-3 and 5-CCAAT-3), which are associated specifically with genes induced at the plasma cell state (35, 36). However, no sequence associated specifically with XBP1 appeared in the final set of *de novo* motifs and so XBP1-associated motif enrichment was not tested amongst PGCNA modules derived from the expression time course.

Finally, assumptions made by cisREAD’s methodology may also limit its utility in predicting certain regulatory relationships. cisREAD employs a correlation-based approach to link cis-regulatory elements to target genes, looking for regulatory elements whose accessibility is a positive or negative predictor of gene expression. Whilst correlation based methods can assign enhancers and silencers to genes to provide biological insight (59–61), they may overlook regulatory relationships where accessibility is not correlated with expression – e.g. priming (62) – or assign spurious relationships where correlation is not indicative of regulation. The construction of sparse LASSO models (selecting the few best predictors), may also struggle to recover transcriptional control by super-enhancers, involving large stretches of individual cis-regulatory elements (63, 64). Whilst the community detection step serves to group together potential co-acting enhancers, this has been done based on both transcription factor footprint and chromatin accessibility. To better capture super-enhancers, at the expense of TF-co-bound units, cisREAD’s community detection step can be performed using only chromatin accessibility. It should also be noted that in a minority of instances, selected regression coefficients reverse sign, with the model coefficient opposite between the correlation between accessibility and expression. This is due to the statistical paradox of suppression, which can arise upon addition of another correlated variable into the model (65). For this reason, we derive the direction of regulation from correlation not coefficient.

Despite these limitations, we show here that application of cisREAD to our *in vitro* B cell differentiation model reveals how a crucial switch from PU.1/SPIB to AP-1 (BATF) reprograms ABC transcription in preparation for plasma cell fate. The overwhelming association with BATF/AP-1 with the ABC state suggests that BATF itself may be responsible for licencing ABC-specific regulatory elements. Observations here, and elsewhere (50), support a role for BATF as a pioneer factor in B cells. Recent experimental work establishing BATFs pioneering function in T cells also supports investigation into BATF pioneering in the B lineage (66). Our work represents a step towards understanding how the B cell cis-regulatory landscape is reshaped during the human immune response.

## DATA AVAILABILITY

ATAC-seq and RNA-seq datasets have been deposited in GEO under the accession GSE219012 (Password/token for reviewers: ehghwggelhgxzmf)

ATAC-seq signal tracks for the 19 samples are viewable on the UCSC genome browser at: https://genome-euro.ucsc.edu/cgi-bin/hgTracks?hgS_doLoadUrl=submit&hgS_loadUrlName=https://raw.githubusercontent.com/AmberEmmett/InVitroBCellsATACSeq/main/InVitroBCellsATACSeq.txt

cisREAD is available as an R package at the GitHub repository https://github.com/AmberEmmett/cisREAD and has been deposited on Figshare at https://figshare.com/articles/software/cisREAD/21821151

Interactive gene co-expression networks are available at https://matthewcare.wixsite.com/pgcna/bcell-detailedtc (Password for reviewers: DlbclJaVu)

## SUPPLEMENTARY DATA

Supplementary Data are available at NAR online.

## Author Contributions

A.E. and A.S. should be regarded as joint first authors and D.W. and R.T as joint senior authors. A.E. designed and carried out data analysis and wrote the initial manuscript. A.S. designed experiments and generated data. M.C contributed to data analysis. G.D. contributed to data generation. R.T. and D.W. conceived and designed the study. All authors read and approved the final draft manuscript. Part of this work was undertaken on ARC4, part of the High Performance Computing facilities at the University of Leeds, UK. The graphical abstract was partially created with BioRender.com.

