## Supplementary Methods and Figures for "Integration of chromatin accessibility and gene expression data with cisREAD reveals a switch from PU.1/SPIB-driven to AP-1-driven gene regulation during B cell activation"

**Transcription factor enrichment in cis-regulatory element clusters**

To investigate differentiation-stage specific transcription factor (TF) binding, k-means clustering of standardized log_2_ normalised chromatin accessibilities was performed for all significant cis-regulatory elements (CREs) predicted to regulate a differentially expressed gene (DEG), with k = 8. Following the identification of cell-stage specific regulatory clusters, TF occupancy enrichment was calculated using a two-sided fisher test for each TF-cluster combination, comparing whether the TF occupancy rate of a cluster is significantly greater than (enriched) or lesser than (depleted) the TF occupancy rate of all other clusters. *De novo* motifs enriched in each cluster were discovered using HOMER findMotifsGenome.pl, compared to a background of non-differentially accessible regions. CREs in each cluster were annotated with HOMER annotatePeaks.pl to derive annotations and nearest gene distance. GC content was calculated using the bedtools nuc command.

**Parsimonious gene co-expression network analysis and TF enrichment**

To explore gene expression dynamics across differentiation, a gene co-expression network was constructed using the Parsimonious Gene Correlation Network Analysis (PGCNA) method (1) . The gene expression data, including 3 additional samples (6h x 2, D6 x1) without matching ATAC-seq, were analysed using DESeq2, identifying 16,296 differentially expressed genes (LRT; BH-FDR 0.01). The VST normalized expression data for the DEG was analysed with PGCNA2 [settings -n 1000, -f1, -b 100] (https://github. com/medmaca/PGCNA/tree/master/PGCNA2) selecting the best clustering using scaled cluster enrichment score. This gave a network with 16,296 nodes and 57,175 edges (available <https://matthewcare.wixsite.com/pgcna/bcell-detailedtc>).

Module names were derived from gene-set over-representation analysis (FDR < 0.1). First a data-set of 18,920 gene signatures was created by merging signatures downloaded from [lymphochip.nih.gov/signaturedb/](https://eur03.safelinks.protection.outlook.com/?url=http%3A%2F%2Flymphochip.nih.gov%2Fsignaturedb%2F&data=05%7C01%7Cbsaem%40leeds.ac.uk%7C9533c00f81d447860a6f08dacd73ebb3%7Cbdeaeda8c81d45ce863e5232a535b7cb%7C1%7C0%7C638048196353428868%7CUnknown%7CTWFpbGZsb3d8eyJWIjoiMC4wLjAwMDAiLCJQIjoiV2luMzIiLCJBTiI6Ik1haWwiLCJXVCI6Mn0%3D%7C3000%7C%7C%7C&sdata=zCOQjN3EuIGOYqHl5Y2oGS%2FUmjyAzvhjEdUyhXoKMy8%3D&reserved=0) (SignatureDB), [www.broadinstitute.org/gsea/msigdb/index.jsp](http://www.broadinstitute.org/gsea/msigdb/index.jsp%20) MSigDB V7.2 (MSigDB C1--C7 and H; excluding C5), Human CORUM complexes with > 2 genes ([http://mips.helmholtz-muenchen.de/corum/#download](https://eur03.safelinks.protection.outlook.com/?url=http%3A%2F%2Fmips.helmholtz-muenchen.de%2Fcorum%2F%23download&data=05%7C01%7Cbsaem%40leeds.ac.uk%7C9533c00f81d447860a6f08dacd73ebb3%7Cbdeaeda8c81d45ce863e5232a535b7cb%7C1%7C0%7C638048196353428868%7CUnknown%7CTWFpbGZsb3d8eyJWIjoiMC4wLjAwMDAiLCJQIjoiV2luMzIiLCJBTiI6Ik1haWwiLCJXVCI6Mn0%3D%7C3000%7C%7C%7C&sdata=8J7BjYaBkYgqWxT3xpMCdOcCBT%2FOFfcvLJmp2X6kV%2F0%3D&reserved=0)), UniProt keywords (parsed XML) and 17 papers (PMIDs:12975453,15550490,19412164,20725040,21179087,23563690,23584089,23584090,23700391,23871637,24138885,24220563,24336226,24644022,25800755,25822800,28735890). A gene ontology gene-set was created using an in-house python script. This parses a gene association file ([geneontology.org/page/download-annotations](https://eur03.safelinks.protection.outlook.com/?url=http%3A%2F%2Fgeneontology.org%2Fpage%2Fdownload-annotations&data=05%7C01%7Cbsaem%40leeds.ac.uk%7C9533c00f81d447860a6f08dacd73ebb3%7Cbdeaeda8c81d45ce863e5232a535b7cb%7C1%7C0%7C638048196353428868%7CUnknown%7CTWFpbGZsb3d8eyJWIjoiMC4wLjAwMDAiLCJQIjoiV2luMzIiLCJBTiI6Ik1haWwiLCJXVCI6Mn0%3D%7C3000%7C%7C%7C&sdata=Jk6irZB2kAg%2BlRoaZEB%2BjXz8N5f5X5f8cXeLsLS%2FT5I%3D&reserved=0) ) to link genes with ontology terms, then uses the ontology structure (.obo file; [purl.obolibrary.org/obo/go.obo](https://eur03.safelinks.protection.outlook.com/?url=http%3A%2F%2Fpurl.obolibrary.org%2Fobo%2Fgo.obo&data=05%7C01%7Cbsaem%40leeds.ac.uk%7C9533c00f81d447860a6f08dacd73ebb3%7Cbdeaeda8c81d45ce863e5232a535b7cb%7C1%7C0%7C638048196353428868%7CUnknown%7CTWFpbGZsb3d8eyJWIjoiMC4wLjAwMDAiLCJQIjoiV2luMzIiLCJBTiI6Ik1haWwiLCJXVCI6Mn0%3D%7C3000%7C%7C%7C&sdata=t9W9%2FXvDvKawmX6Rhu3b1HDAwH34ZzVS%2FqYAlMapDvk%3D&reserved=0) ) to propagate these terms up to the root. The resultant gene-set contained 22,891 terms. The gene-ontology and gene-signatures sets were merged to give a final signature set of 41,811 terms. Enrichment of modules for signatures was assessed using a hypergeometric test, where the draw is the module genes, the successes are the signature genes and the population is the genes present on the platform.

Transcription factors were then linked to PGCNA modules by identifying TF footprints in significant cis-regulatory elements selected to regulate expression of genes in each module. TF enrichment was then calculated using a two-sided fisher test, comparing TF occupancy in CREs linked to that module to the occupancy of CREs not linked to that module.

**SPIB, PU.1 and BATF ChIP-seq**

ChIP-seq targeting PU.1 and BATF in GM12878 was downloaded from ENCODE (IDR-threshold peaks) to obtain transcription factor binding sites (2). SPIB binding sites were obtained from the union of ChIP-seq peaks called in OCILy3 and OCILy10 (macs2 q value < 0.01) using data from Care et al. 2014, realigned to hg38 and processed as described (3). Binding sites for each factor were intersected (using bedtools intersect) with differentially accessible regions (DARs).

**Clustering of PU.1/SPIB and AP-1 target genes**

Genes with cisREAD-linked regulatory elements with PU.1/SPIB and/or AP-1 footprints, whose accessibility was positively correlated with expression, were defined as potential PU.1/SPIB and/or AP-1 targets. These genes were then k-means clustered by expression (k=5, incremented until early and late ABC clusters were separated). Enrichment of ChIP-seq transcription factor binding sites (for PU.1, SPIB and BATF) was calculated for cis-regulatory elements with PU.1/SPIB and/or AP-1 footprints linked to each of the 5 expression clusters. This was done using a one-sided fisher test, comparing occupancy of a TF (PU.1, SPIB or BATF) in CREs with a given footprint (PU.1/SPIB, PU.1/SPIB + AP-1, AP1) linked to one of the five clusters, to occupancy of the same TF in all DARs without the given footprint.

Genes in each expression cluster, linked to either PU.1/SPIB, PU.1/SPIB + AP-1, or AP-1 footprints, were tested for enrichment of Gene Ontology (GO) biological processes (4, 5), Kyoto Encyclopaedia of Genes and Genomes (KEGG) pathways (6), Hallmark Molecular Signature DataBase (MSigDB) gene signatures (7) and Staudt lab gene signatures (8). Enrichment was first calculated using a one-sided fisher test, to test gene-set over-representation in TF targets in a cluster, compared to all other clusters (targeted by any of the 3 TFs.) Gene sets enriched with BH-adjusted p < 0.1 were considered significantly enriched. This was used to define potential ‘temporally regulated’ gene sets. Enrichment was also performed against genes with a similar expression profile without the given TF footprint(s) to identify gene sets preferentially regulated by a TF at a given differentiation stage (‘TF-specific regulated’).

To obtain background sets for gene set over-representation, similarly expressed genes were identified by training an XGBoost classifier (using the ‘XGBoost’ R package) on the five expression clusters and predicting the cluster label for all other differentially expressed genes, only linked to CREs without the TF footprint(s). An XGBoost model, using the ‘multi:softprob’ objective function, was trained by five-fold cross validation (stratified folds), to identify the number of decision trees resulting in the lowest mean multi-classification error (nrounds = 47.) Genes were assigned a cluster label if the XGBoost-predicted probability of belonging to a class exceeded 0.9.

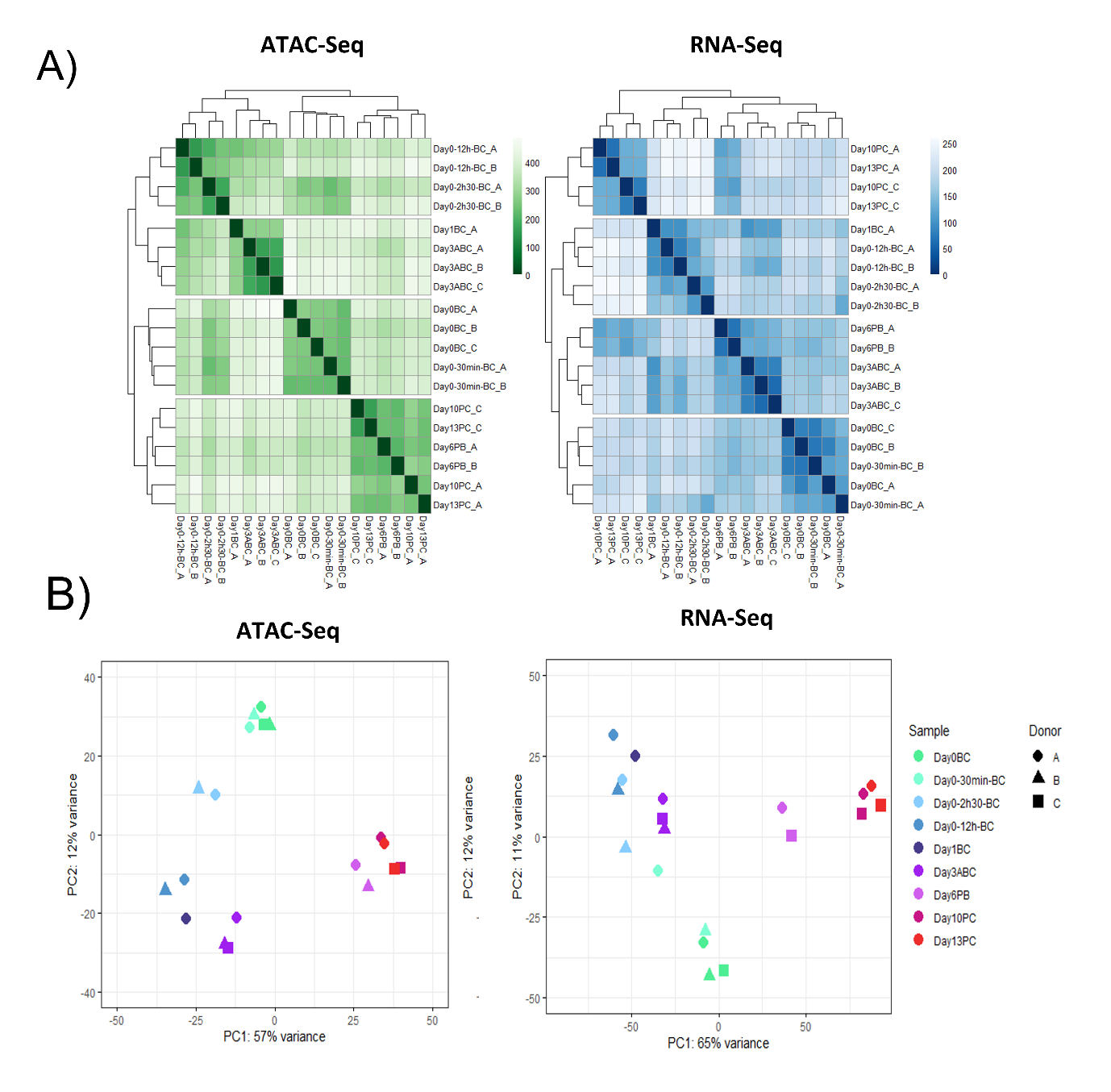

Figure S1. A) Hierarchical clustering of Euclidean sample distances for ATAC-seq and RNA-seq datasets, calculated from VST normalized counts. B) Principal components analysis biplot for ATAC-seq and RNA-seq datasets, showing PC1 and PC2 calculated from VST normalised counts.

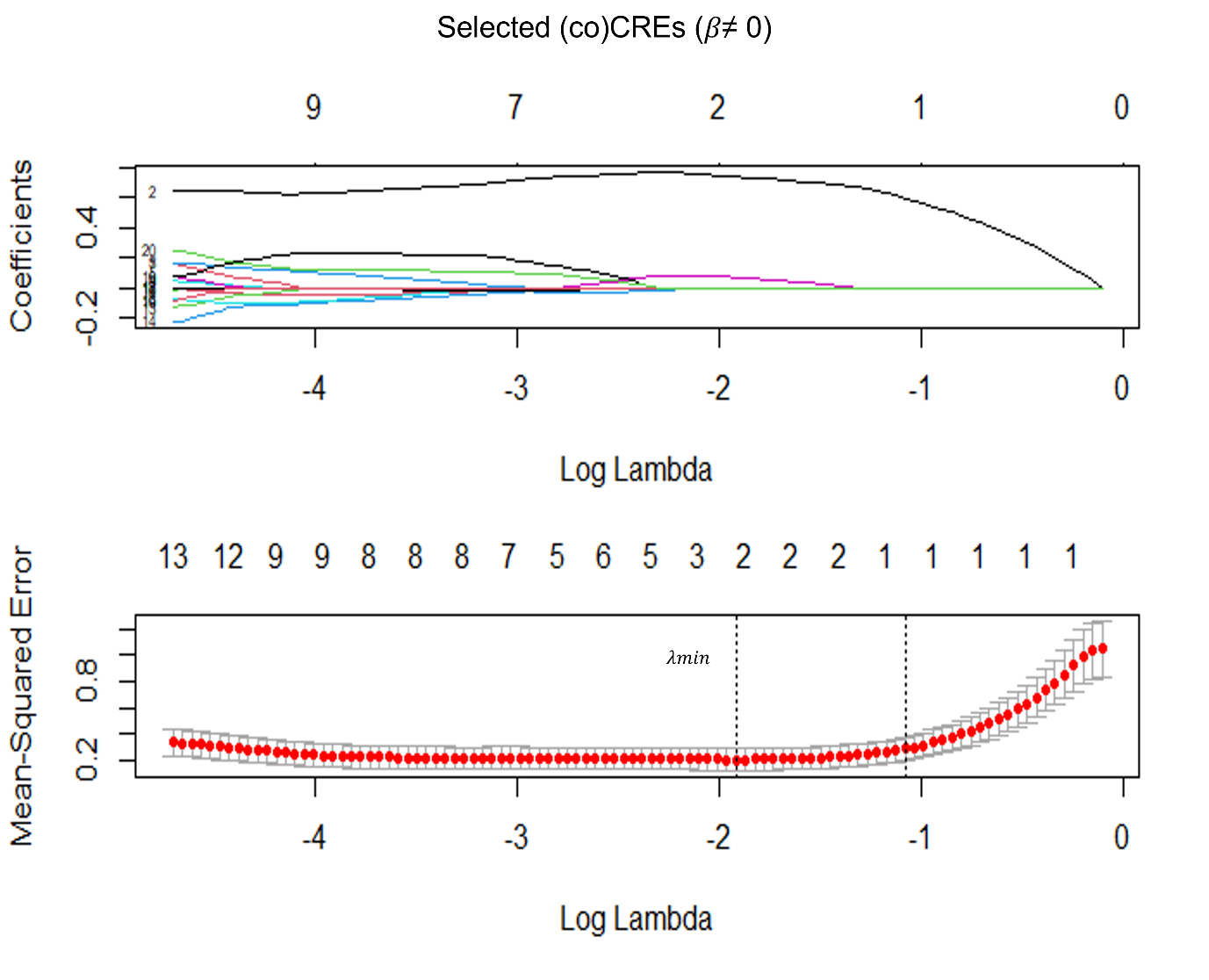

Figure S2. Model selection for BLIMP1 gene, the optimum value of lambda (LASSO tuning parameter) is chosen as that with the minimum mean squared error (lambda-min) during cross-validation, two (co)CRE predictors have non-zero coefficients in the optimum model.

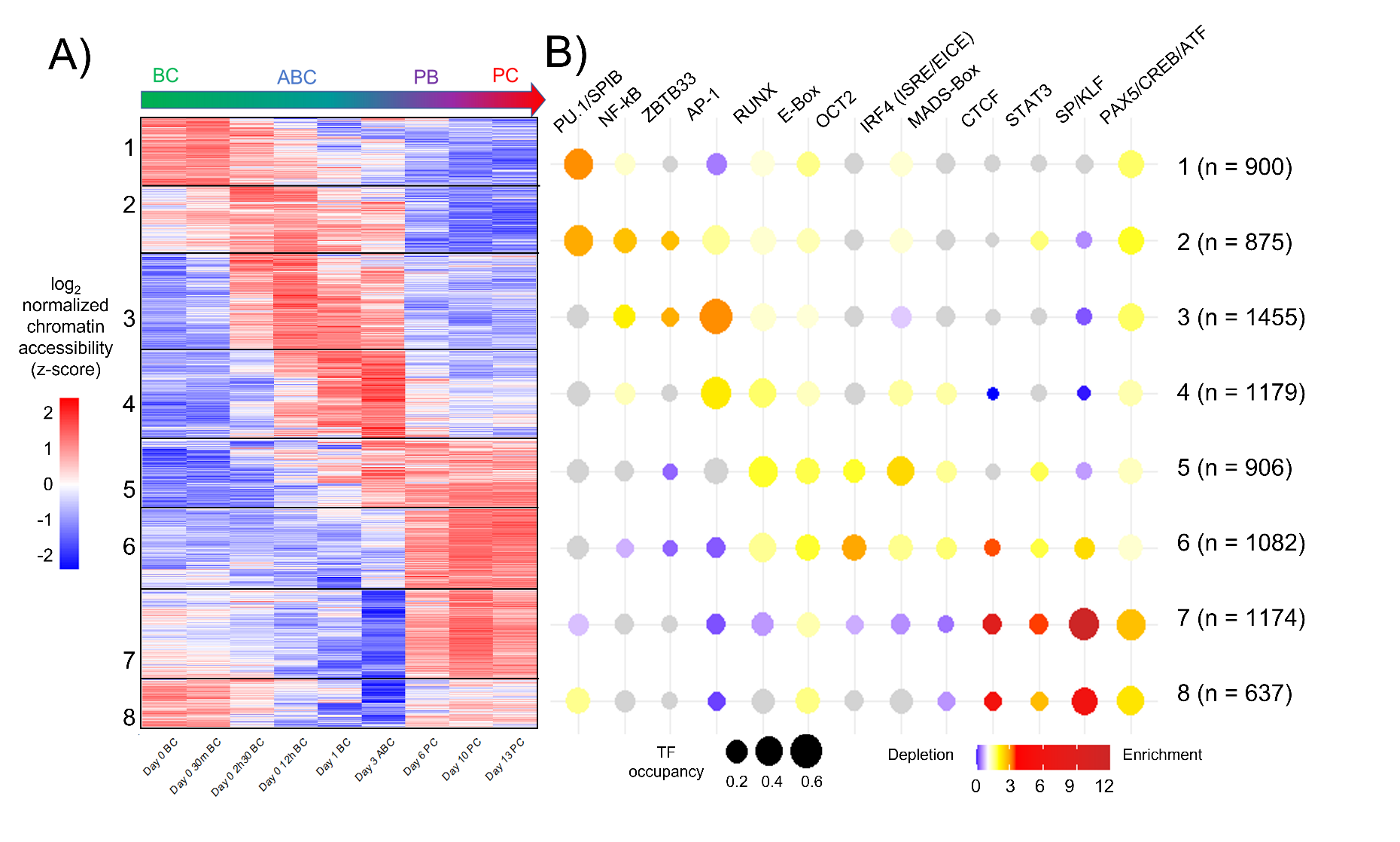

Figure S3. TF footprint enrichment in CREs significantly linked to gene upregulation, derived from BMO inputs to cisREAD. A) Heatmap showing log_2_ normalized chromatin accessibility (z-score) of cis-regulatory elements significantly linked to gene expression, k-means clustered (k = 6) by z-score chromatin accessibility. B) Bubbleplot showing enrichment of TF occupancy in each cluster. Size of bubbles gives the proportion of each cluster harbouring a TF footprint, colour shows significant (p < 0.05, two-sided Fisher test) enrichment (fold-change between cluster and other clusters > 1, red) or depletion (fold-change between cluster and other clusters < 1, blue), grey represents no significant enrichment. N gives the number of CREs in each cluster.

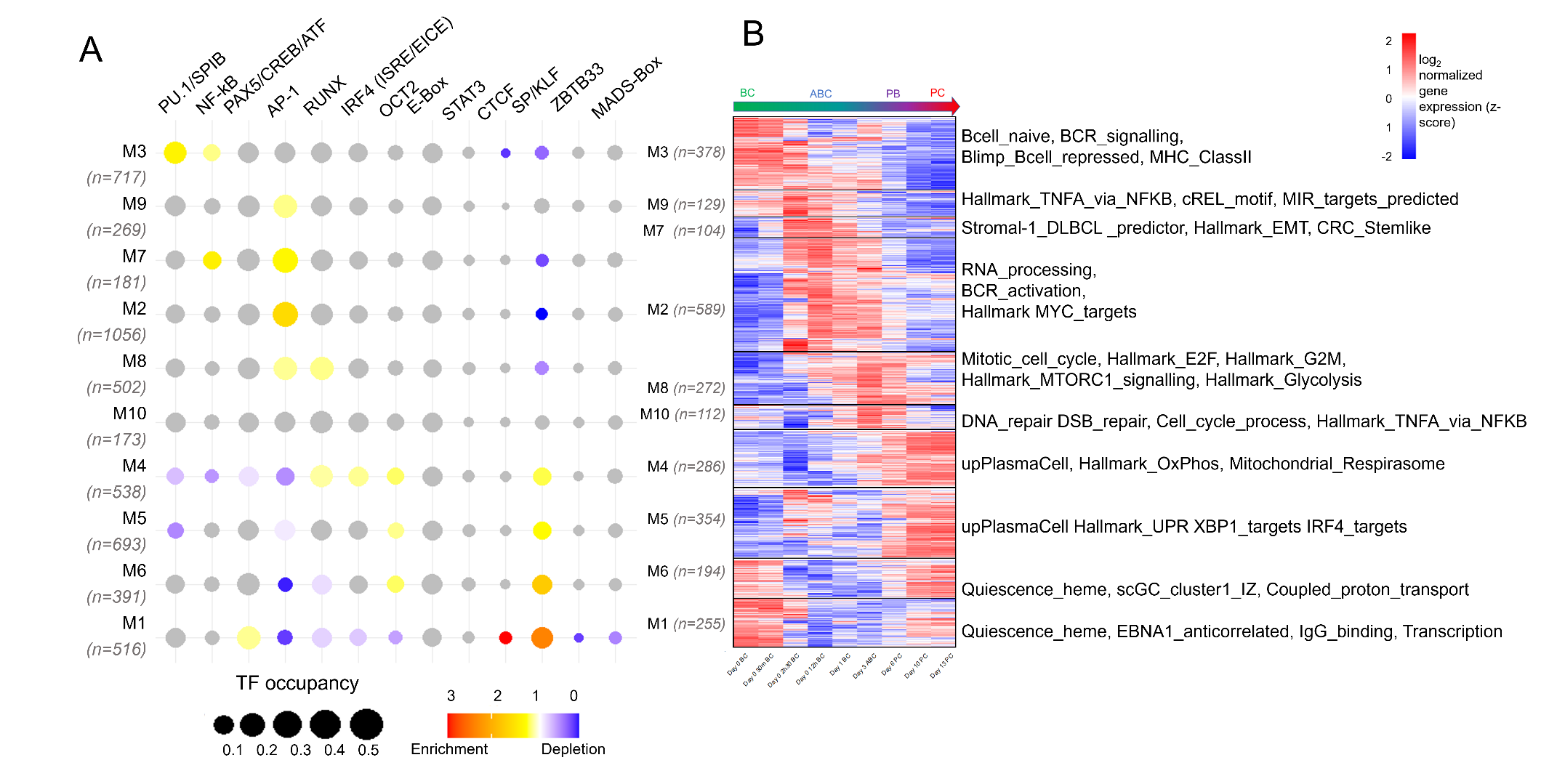

Figure S4. TF footprint enrichment in CREs significantly linked to upregulation of PGCNA modules, derived from BMO inputs to cisREAD. A) Bubbleplot showing enrichment of TF occupancy in each cluster. Size of bubbles gives the proportion of each cluster harbouring a TF footprint, colour shows significant (p < 0.05, two-sided Fisher test) enrichment (fold-change between genes in module and genes not in module > 1, red) or depletion (fold-change < 1, blue), grey represents no significant enrichment. N gives the number of genes with significant CREs in each module. B) Heatmap showing log_2_ normalized gene expression (z-score) of genes with significantly linked CREs, module names reflect enriched gene sets in each module.

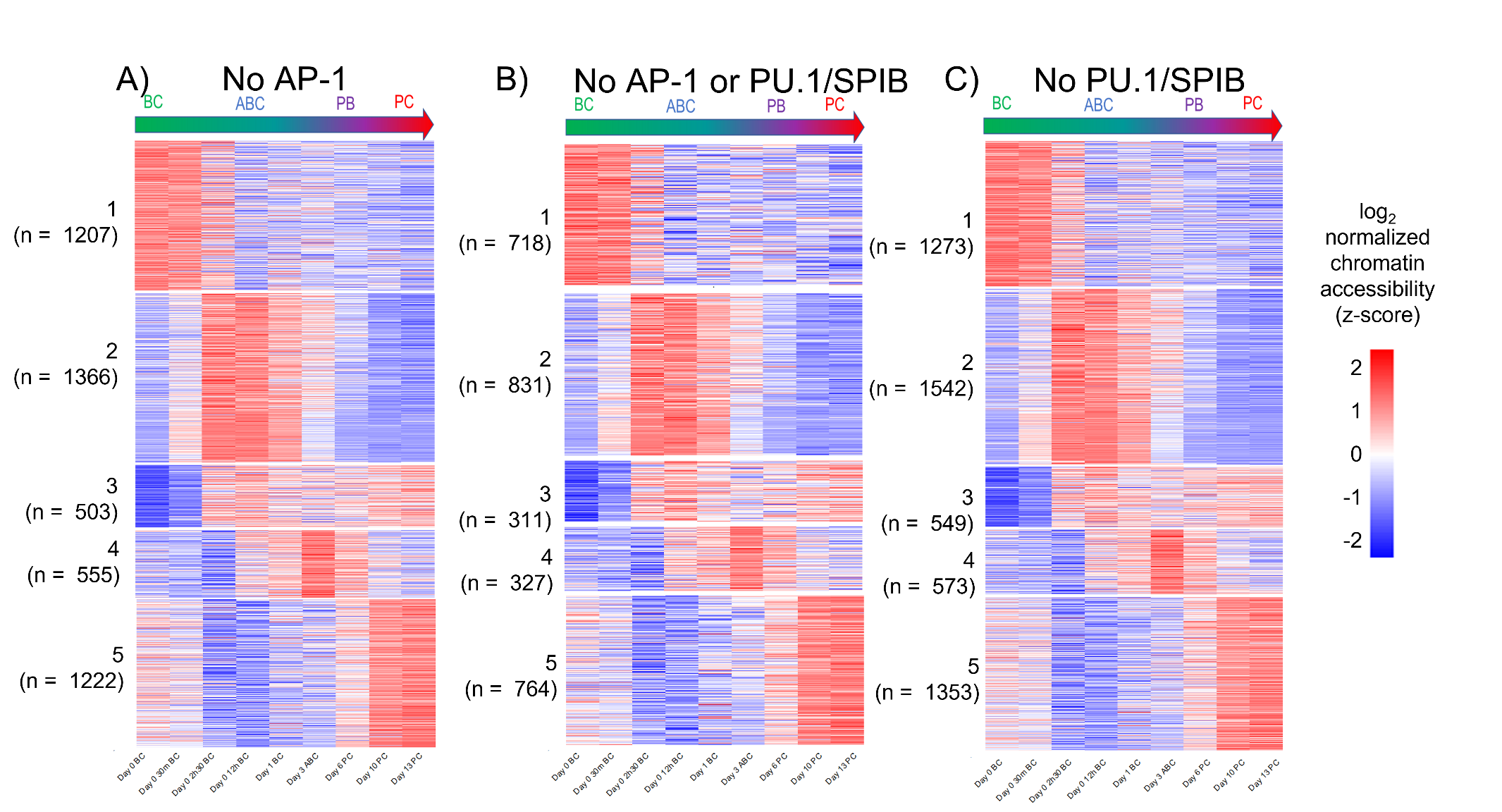

Figure S5. Background genes for gene set over-representation analysis of PU.1/SPIB and/or AP-1 target gene clusters, against similarly expressed genes not linked to the factor(s). For each analysis, similarly expressed genes were obtained by training a machine learning classifier on five expression clusters, and predicting the cluster label for all other differentially expressed genes not linked to CREs with given footprint(s). N gives number of genes, not linked to the factor(s) predicted to belong to each cluster. Heatmaps show z-score log_2_ normalized gene expression for: A) Similar expressed genes not linked to AP-1, B) similarly expressed genes not linked AP-1 or PU.1/SPIB, C) similarly expressed genes not linked to PU.1/SPIB.
